# Basal spot junctions of epithelial tissues respond to morphogenetic forces and regulate Hippo signaling

**DOI:** 10.1101/2023.09.17.558144

**Authors:** Benjamin Kroeger, Samuel A. Manning, Yoshana Fonseka, Viola Oorschot, Simon A. Crawford, Georg Ramm, Kieran F. Harvey

## Abstract

Organ size is controlled by numerous factors including mechanical forces, which are mediated in part by the Hippo pathway. In growing *Drosophila melanogaster* epithelial tissues, cytoskeletal tension influences Hippo signalling by modulating the subcellular localisation of key pathway proteins in different apical domains, namely adherens junctions, the sub-apical region and the medial apical cortex. Here, using both electron and light microscopy, we have discovered the existence of basal spot junctions in *D. melanogaster* epithelial tissues, and that they respond to morphogenetic forces and also influence Hippo signalling. Like adherens junctions, the Warts kinase is recruited to basal spot junctions via Ajuba and E-cadherin, which prevent Warts activation by segregating it from upstream Hippo pathway proteins. Basal spot junctions are prominent when tissues undergo morphogenesis and are highly sensitive to fluctuations in cytoskeletal tension. Basal spot junctions are distinct from focal adhesions, but the latter profoundly influences the abundance of spot junctions by modulating the basal-medial actomyosin network and tension experienced by spot junctions. Thus, basal spot junctions potentially couple morphogenetic forces to Hippo pathway activity and organ growth.

## INTRODUCTION

Signalling pathways are used throughout life to allow organisms to detect different stimuli and control appropriate behavioural responses. Signalling pathways transduce information on different scales, e.g., humoral signals that allow communication between independent organs, signals within organs from non-neighbouring cells, and short-range signals between neighbouring cells or within the same cell. The majority of signalling pathways are regulated by diffusible ligands that engage membrane-bound receptor proteins, whilst neighbouring cells can also communicate via transmembrane proteins, e.g. cadherins ^1^. In polarized cells, like epithelial cells, most signalling events have been described to occur at the apical cell surface. For example, ligand-receptor interactions are thought to occur predominantly at apical membranes and often lead to receptor internalisation in vesicles ^1^. Cell-cell signalling events have also been described as occurring primarily in the apical regions of epithelial cells. For example, in *Drosophila melanogaster* imaginal discs, the atypical cadherins Fat and Dachsous localise to the sub-apical region of epithelial cell-cell contacts and form a physical complex to signal between neighbouring cells ^2^. In addition, major mechanical forces generated by cell-cell adhesion and cytoskeletal tension are mediated by E-cadherin (E-cad) complexes, which are enriched at adherens junctions (AJs), and are apically located ^3^. By contrast, far fewer signalling events have been described to occur at the basal regions of epithelial cells. One notable exception is integrin-mediated signalling; basally located integrins contact the extracellular matrix (ECM) and facilitate cell signalling, as well as adhesion ^4,5^.

A signalling pathway that is not principally regulated by diffusible ligands and their cognate transmembrane receptors is the Hippo pathway. This complex and highly conserved signalling network, which controls organ size and cell fate and is deregulated in many human cancers, responds to mechanical forces and multiple cell biological cues such as adhesion and polarity ^6–11^. In *Drosophila melanogaster* wing imaginal discs, which are epithelial tissues, apical cell membranes and apical cell junctions have been identified as key sites for Hippo signalling regulation ^12–14^. Increased cytoskeletal tension, regulated by actomyosin contractility, promotes activity of the Yorkie (Yki) transcriptional co-activator protein and wing growth ^12^. Mechanistically, increased actomyosin contractility recruits the mechanosensitive protein Ajuba (Jub) and the central Hippo pathway kinase Warts (Wts) to AJs, where Jub is thought to sequester Wts in an inactive state ^12^. Juxtaposed to AJs on their apical side are the sub-apical regions of epithelial cells, where many Hippo pathway proteins can localise to. These include Kibra, Expanded (Ex) and Merlin (Mer), which recruit the Hippo pathway core kinase cassette [the Hippo (Hpo) and Wts kinases and the Salvador (Sav) and Mats adaptor proteins], and catalyse the activation of Hpo and Wts ^13^. Kibra and Mer can also recruit the core kinase cassette to the medial apical cortex and catalyse activation of both Hpo and Wts^14^. Ultimately, Wts phosphorylates Yki ^15,16^, which dictates the rate at which it shuttles between the nucleus and cytoplasm ^17^. Yki controls transcription by forming a physical complex with the Scalloped (Sd) DNA binding protein ^18–20^.

Currently, it is not clear if the sub-apical region and medial apical cortex of epithelial cells are the only sites of Hippo pathway regulation or whether other subcellular regions are also important. To investigate this, we employed electron microscopy and fluorescence microscopy to detect the subcellular localisation of endogenously tagged Hippo pathway proteins in vivo. In doing so, we discovered a previously uncharacterized pool of both the Wts and Jub proteins at the basal-most region of the lateral membranes. We revealed these regions to be AJ-like spot junctions and found that they were prominent in multiple *D. melanogaster* epithelial tissues, when undergoing morphogenesis. As with the AJ-associated Jub-Wts pool, basal Jub-Wts accumulated at E-cad rich sites, whilst other core Hippo pathway proteins were not observable in basal spot junctions. The basal spot junction pool of Wts was highly sensitive to fluctuations in cytoskeletal tension and therefore might be important for regulating Hippo pathway activity in response to mechanical forces exerted during tissue morphogenesis.

## RESULTS

### Detection of endogenous proteins in vivo by electron microscopy

Defining the localisation of proteins within cells can reveal insights into their function and their relationships with other proteins. Studies of Hippo pathway proteins in imaginal discs using antibodies, endogenously tagged proteins and transgene-encoded proteins have defined AJs, the sub-apical region and the medial apical cortex as important subcellular regions of Hippo pathway regulation. To explore Hippo pathway protein subcellular localisation with nanometre resolution, we employed electron microscopy. Given a paucity of high-quality antibodies for Hippo pathway proteins, we used CRISPR-Cas9 genome editing to generate a suite of *D. melanogaster* strains where different Hippo pathway genes were tagged at their endogenous loci with fluorescent proteins ^17,21–23^. To improve our ability to detect Hippo pathway protein localisation by electron microscopy, we adopted a system developed in zebrafish ^24^, where a GFP-Binding Peptide (GBP) ^25^ is fused to an engineered version of the soy bean enzyme, APEX ^26^, which converts diffusive 3,3′-diaminobenzidine (DAB) into an insoluble osmiophilic polymer in the presence of H_2_O_2_. When exposed to OsO_4,_ this polymer is converted into an electron-dense precipitate (Figure S1A). We generated transgenic *D. melanogaster* that express APEX2-GBP under the control of the UAS promoter, which affords spatial and temporal control of transgene expression. *D. melanogaster* strains were generated that were homozygous for the fluorescently-tagged protein of interest and expressed APEX2-GBP under the control of *en-GAL4*. The successful expression of APEX was confirmed with a DAB reaction, with the colour reaction evident specifically in the posterior compartment of the wing imaginal disc, where *en-GAL4* is active (Figure S1B). Importantly, the ability to spatially control the expression of APEX2-GBP with the UAS-GAL4 system provides an internal control that allows better determination of signal to noise by comparing neighbouring cells at the anterior-posterior compartment boundary, only one of which expresses APEX2-GBP (Figure S1B).

### The Warts kinase localises to basal puncta on lateral membranes of epithelial cells

Initially, we used electron microscopy to assess the subcellular localisation of Wts, given it is the key downstream kinase in the Hippo pathway and has a relatively specific localisation pattern at the AJs in the apical region of imaginal disc cells ^12,13,27^. First, using confocal microscopy, we assessed whether Wts-Venus localisation and growth control function was affected by expression of APEX2-GBP. Wts-Venus was clearly detectable at AJs in both the control anterior compartment of wing imaginal discs, as well as the posterior compartment, which expressed APEX2-GBP, indicating that it did not affect Wts-Venus localisation (Figures S1C and S1C’). APEX2-GBP also did not substantially impinge on Wts function, as wing size was only slightly reduced in *wts-Venus* animals that expressed APEX2-GBP (Figures S1D-S1G).

An electron-dense precipitate, reflecting Wts-Venus was observable at apical cell-cell junctions of wing disc cells, which are consistent with AJs (Figures 1A, 1B and S1H-S1H’’). In *wts-Venus* animals that expressed APEX2-GBP under the control of *en-GAL4*, we were able to compare cross sections of cells at either side of the anterior-posterior boundary and detect a Wts-Venus signal at the AJ of an APEX2-GBP-expressing cell and the absence of such a signal in the juxtaposed control cell (Figures 1A and S1H’). Wts-Venus protein was also observed in the sub-apical region, above the AJs (Figure S1H’’). To our surprise, a pool of Wts-Venus was also observable by electron microscopy at the basal-most region of the lateral cell membranes (Figures 1C and S1I-1I’’). To our knowledge, a basal pool of Wts protein, or any other core Hippo pathway protein, has never before been reported in *D. melanogaster* epithelial cells. Similarly, we are unaware of reports of core Hippo pathway proteins localising to the basal region of cells in vivo in other species.

**Figure 1.**
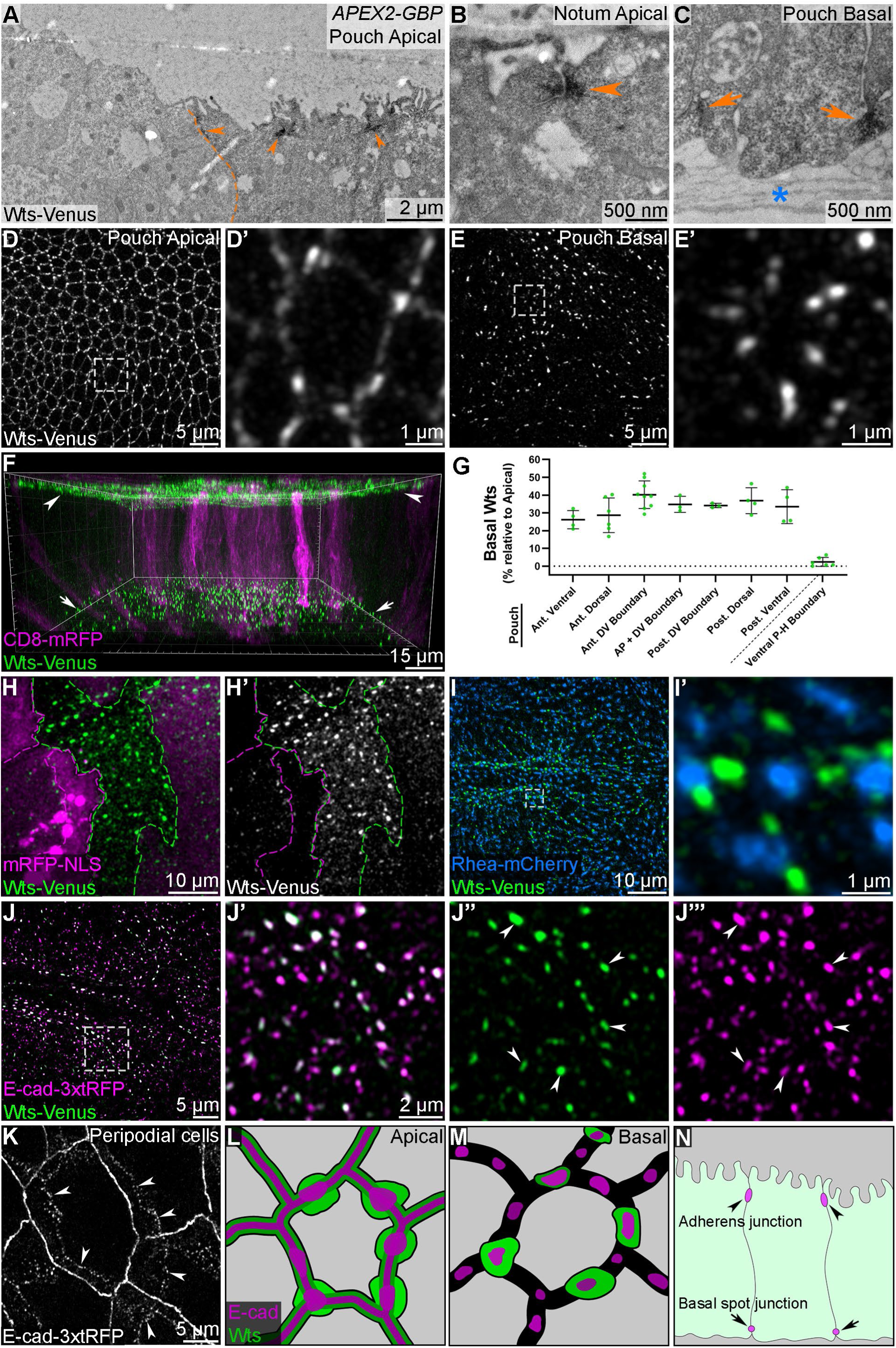
Identification of a basolateral punctate pool of Warts protein in cells of *Drosophila* epithelial tissues. **(A-C)** Electron micrographs of *D. melanogaster* third instar larval wing imaginal discs from *wts-Venus* animals that also express APEX2-GBP in the posterior compartment, under the control of *en-GAL4*. A dashed orange line indicates the anterior-posterior compartment boundary in (**A**). Note the elevated DAB signal at the apical junctions (orange arrowheads) and in the basal-most region of the lateral membranes (orange arrows). Elevated DAB signals were present specifically in cells of the posterior wing disc, but not in control, anterior cells. The basement membrane is marked with a blue ‘*’ in (**C**). **(D-E’)** Super-resolution confocal microscope images of either the apical or basal regions of third instar larval wing imaginal discs. Wts-Venus is grayscale, boxed regions are shown at higher magnification. **(F)** Three-dimensional reconstruction image of Wts-Venus (green) in a wing imaginal disc harbouring small CD8-mRFP+ clones (magenta). Wts-Venus is evident in both the apical (arrowheads) and basal-most (arrows) regions of the tissue. **(G)** Chart displaying quantification of basal relative to apical junctional Wts-Venus protein in small clones from different regions of wing imaginal discs, imaged as in (**F**). n ≥ 3 clones for each subregion of the pouch, and n = 6 clones in the ventral pouch-hinge boundary fold. **(H-H’)** Confocal microscope image of the basal region of a wing imaginal disc. Wts-Venus is green and mRFP-NLS is magenta in the merged image (**H**) and Wts-Venus is greyscale in the non-merged image (**H’**). This genetically mosaic tissue is comprised of three cellular populations: *wts-Venus* homozygous clones (expressing Wts-Venus, green), *mRFP-NLS* homozygous twin-spot clones (magenta), and heterozygous cells (expressing both Wts-Venus, green and mRFP-NLS, magenta) Dashed lines highlight homozygous clone boundaries. **(I-I’)** Confocal microscope images of the basal region of a wing imaginal disc. Wts-Venus is green and Rhea-mCherry is blue. The boxed region is shown at higher magnification in (**I’**). **(J-J’’’)** Confocal microscope images of overlapping basal Wts-Venus (green) and E-cad-3xtRFP (magenta) puncta are indicated (arrowheads in **J’’ and J’’’**). The boxed region in (**J**) is shown at higher magnification in (**J’-J’’’**). (K) Confocal microscope image of E-cad-3xtRFP in the peripodial epithelium of a wing imaginal disc. Note the discontinuous basal E-cad puncta in peripodial cells (arrowheads). **(L-N)** Schematic representations of apical and basal Wts and E-cad localization. Apical cell junctions have continuous, belt-like AJs associated with Wts protein, whilst basal cell-cell interfaces have discontinuous E-cad puncta, some of which are associated with Wts puncta. Scale bars are indicated in image panels. All confocal microscopy images are maximum intensity projections except for (**J-J’’’**), which are single z-slices. Data is representative for at least 10 specimens, except for (**A-C**), where 3 wing imaginal discs were examined. See also Figures S1 and S2.

To confirm the presence of basal Wts in epithelial cells, we performed light microscopy studies on imaginal discs from *wts-Venus* animals. Wts-Venus was enriched at AJs, consistent with published reports of independent *D. melanogaster* strains expressing Wts proteins tagged with fluorescent proteins or other epitopes ^12,13,27^, and also in basal puncta, thus corroborating our electron microscopy studies (Figures 1D-1E’). Basal Wts puncta were not evenly distributed throughout wing discs, but instead were prominent in specific regions of these tissues, such as the wing pouch (Figure S2B). To better visualize this, we generated high resolution 3-D reconstructions of larval wing imaginal discs, which enabled visualization of Wts-Venus on both the apical and basal surfaces (Figure 1F and Video S1). This indicated that the majority of Wts localizes to either the apical or basal cell membranes, with minimal protein in the cytoplasm. By generating small RFP-positive clones, we were able to quantify the relative amounts of apical and basal Wts. This revealed that the apical vs basal ratio of Wts varied throughout different regions of wing discs and that basal Wts was more variable across wing imaginal discs than AJ-associated Wts (Figure 1G). As a final confirmation that the observed basal Wts-Venus signal did not reflect any form of non-specific fluorescence, we recombined *wts-venus* onto an FRT82B chromosome and generated clones of *wts-venus* and RFP-expressing control tissue using mitotic recombination. Clones expressing Wts-Venus displayed basal punctate Wts and apical junctional Wts, whilst control RFP clones did not (Figures 1H-1H’ and S2A-S2A’’’), thus validating the existence of a basal pool of Wts protein. Further, basal Wts puncta were observed using independent *D. melanogaster* strains expressing different tagged Wts proteins – Wts-GFP^12^, and mCitrine-Wts-mTFP ^27^ (Figures S2D-S2E), and therefore is not specific to the *wts-Venus* strain generated by our laboratory. To determine whether basal Wts puncta are specific to wing imaginal discs or are a more general phenomenon, we assessed third instar larval eye-antennal discs, leg discs and halteres. In each tissue, Wts was evident in basal puncta, indicating that the localization of Wts to basal puncta is a general feature of third instar larval imaginal discs (Figures S2F-S2G and data not shown).

Integrins localise to the basal medial region of epithelial cells, where they nucleate focal adhesion complexes that anchor cells to the ECM ^4,5^. Further, Integrins and associated signaling proteins like SRC can regulate Hippo pathway activity in mammals ^28–32^, although this regulatory link is not conserved in *D. melanogaster* ^33,34^. To determine whether Wts resides in basal focal adhesions, we assessed the subcellular localisation of Wts and Rhea (known as Talin in mammals), using a *D. melanogaster* strain expressing endogenously tagged versions of each protein (Wts-Venus and Rhea-mCherry). Both Wts and Rhea were evident in the basal regions of wing imaginal disc cells, although Rhea was more broadly visible throughout the wing disc compared to Wts, which was present in basal puncta predominantly in the wing pouch and sub-regions of the hinge (Figures 1I). However, Wts and Rhea did not obviously co-localise; instead, their localisation was largely mutually exclusive at the basal plane of wing imaginal disc cells (Figure 1I’). Rhea appeared to be localised to the basal medial region of disc epithelial cells, while Warts was juxtaposed to Rhea and present on the basal-most part of the lateral membranes (Figures 1C and 1I-1I’). A similar distribution of Wts and Rhea was also evident in larval eye discs, where basal Wts was most evident posterior to the morphogenetic furrow (Figures S2G-S2H). Therefore, basal Wts does not localise to integrin-based focal adhesions but is juxtaposed to them in a similar focal plane.

### E-cadherin forms spot adherens junctions in the basal-most domain of epithelial cell lateral membranes

In growing third instar larval wing imaginal discs, Wts is enriched at AJs, which mediate adhesion between neighbouring epithelial cells, and also connect to the actomyosin cytoskeleton ^3^. Several proteins are enriched at AJs, including E-cad, α-catenin and β-catenin ^3^. E-cad concentrates at AJs in *D. melanogaster* imaginal discs and has been documented at lower levels in vesicles and on basolateral membranes ^35^, but, to our knowledge, not in basal puncta. Given that Wts was not observable in focal adhesions, we investigated other adhesion complex proteins that it might co-localise with and focussed on AJ proteins. Interestingly, as well as being observable at AJs, E-cad-3xtRFP was also evident in distinct puncta in the basal-most region of lateral cell membranes but not prominently at other sites along the lateral membranes. E-cad strongly co-localised with Wts at both basal puncta and at AJs (Figures 1J-1J’’’ and S2C-S2C’’’). E-cad’s localisation both apically and basally was particularly evident in the squamous cells of the peripodial epithelium of larval wing discs, where it was continuously distributed around the apical AJs, and present in regularly spaced puncta at the basal-most part of the lateral membranes (Figure 1K). Close spatial analysis of these E-cad foci revealed a high degree of co-localisation with Wts (Figures S2I-S2J’, schematised in Figures 1L-1N).

At AJs, E-cad ligates with E-cad in neighbouring cells and in this way mediates cell-cell adhesion ^3^. To determine whether the E-cad that we observed in basal puncta on lateral membranes is also likely to interact with E-cad on neighbouring cells and mediate cell-cell adhesion we generated a *D. melanogaster* strain that expressed E-cad fused to one of two different fluorescent proteins (E-cad-GFP and E-cad-3xtRFP). We then generated neighbouring clones that expressed either one or the other E-cad fluorescent protein. By focussing on clone boundaries, we observed close apposition of E-cad at apical AJs of neighbouring cells (Figures 2A-2B). Likewise, E-cad in basal puncta always associated with apposed E-cad puncta of neighbouring cells (Figures 2C-2D). Further, basal E-cad puncta strongly co-localised with α-catenin and were associated with basal-medial F-actin bundles (Figures 2E-2G’’’). Finally, RNAi-mediated depletion of E-cad in small wing imaginal disc clones caused E-cad to be lost from the apposing membranes of neighbouring cells, both at AJs and in basal puncta (Figures 2H-2I’). Collectively, this indicates that these basal puncta, which form at the basal-most point of the lateral membranes, are an AJ-like complex that is associated with the cytoskeleton, and that they likely facilitate cell-cell adhesion. As such, we termed these structures basal spot junctions.

**Figure 2.**
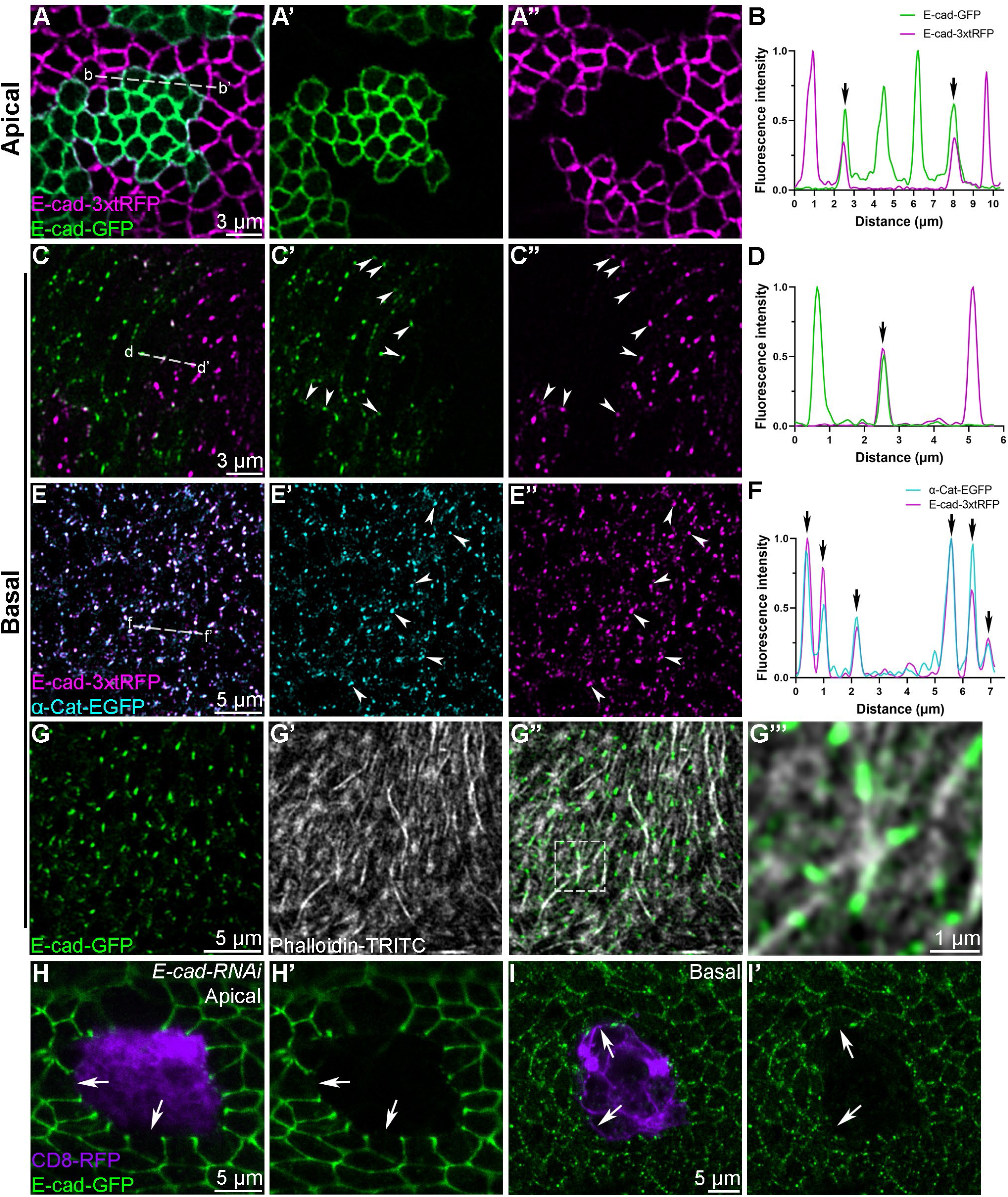
*Drosophila* epithelial tissues possess punctate adherens junction-like structures at the basal-most region of the lateral membranes. (**A, C and E**>) Super-resolution Airyscan images of either the apical (**A-A’’**) or basal (**C-C’’ and E-E’’**) regions of third instar larval wing imaginal discs. E-cad-3xtRFP is magenta, E-cad-GFP is green and α-Cat-EGFP is cyan. Dashed lines in the merged images indicate the regions quantified in (**B, D and F**). Note the basal E-cad puncta at clone boundaries (white arrowheads in **C’ and C’’**) and the overlapping basal E-cad and α-Cat puncta (white arrowheads in **E’ and E’’**). **(B, D and F)** Line profile analyses of fluorescence intensity of E-cad (**B and D**) and of α-Cat and E-cad (**F**) across the cell junctions at clone boundaries. The line profiles run from left to right, i.e., b-b’ in (**A’**) is graphed in (**B**). Spatial correlation at cell boundaries of apposed E-cad-GFP and E-cad-3xtRFP cells at both apical and basal regions is indicated (black arrows in **B and D**), as is the overlap of basal E-cad and α-Cat puncta (black arrows in **F)**. **(G-G’’’)** Super-resolution Airyscan images of the basal hinge region of third instar larval wing imaginal discs. E-cad-GFP is green and Phalloidin-TRITC (marks F-actin) is in grayscale. The boxed region is shown at higher magnification in (**G’’’**). **(H and I)** Confocal microscope images of the apical and basal regions of third instar larval wing imaginal discs harboring E-cad-RNAi-expressing clones (marked with CD8-FRP, purple). E-cad-GFP (green) is depleted in the clones and is also absent from the membranes of directly juxtaposed wild-type cells, both apically (**H and H’**) and basally (**I and I’**) (white arrows). Scale bars are indicated in image panels. All images are single z-slices. Data is representative for at least 5 specimens. See also Figure S2.

### Warts localises to basal spot junctions via Ajuba and E-cadherin

In wing imaginal discs, Wts is recruited to AJs by Jub in a cytoskeletal tension-dependent manner, where it is held in an inactive complex ^12,27^. In this way, the Hippo pathway is thought to be regulated by mechanical forces in order to contribute to organ size control ^12^. Strikingly, we observed that basal spot junction-associated Wts-Venus strongly co-localised with Jub-mKate2, as it does at AJs (Figures 3A-3B). Jub-GFP was also readily observable in basal spot junctions in both larval eye and antennal discs, where it co-localised with E-cad-3xtRFP (Figures 3C-3D and data not shown). Given that Jub is required for Wts localisation at AJs ^12^, we tested if the same was true for basal spot junction-associated Wts. Indeed, RNAi-mediated depletion of Jub strongly reduced the association of Wts at both basal spot junctions and AJs (Figures 3E-3F’ and S3A-S3C’). Notably, apical junctional Wts was reduced, but still present, upon Jub depletion, while it was almost undetectable at basal spot junctions (Figures 3E-3H and S3B-S3C’). To validate these results, we found that both jub-RNAi lines caused a strong depletion in Jub-GFP levels (Figure S3D-E). This indicates that Jub is essential for Wts to accumulate at basal spot junctions and that Wts association with AJs is less reliant on Jub, possibly because it also associates with proteins other than Jub at AJs.

**Figure 3.**
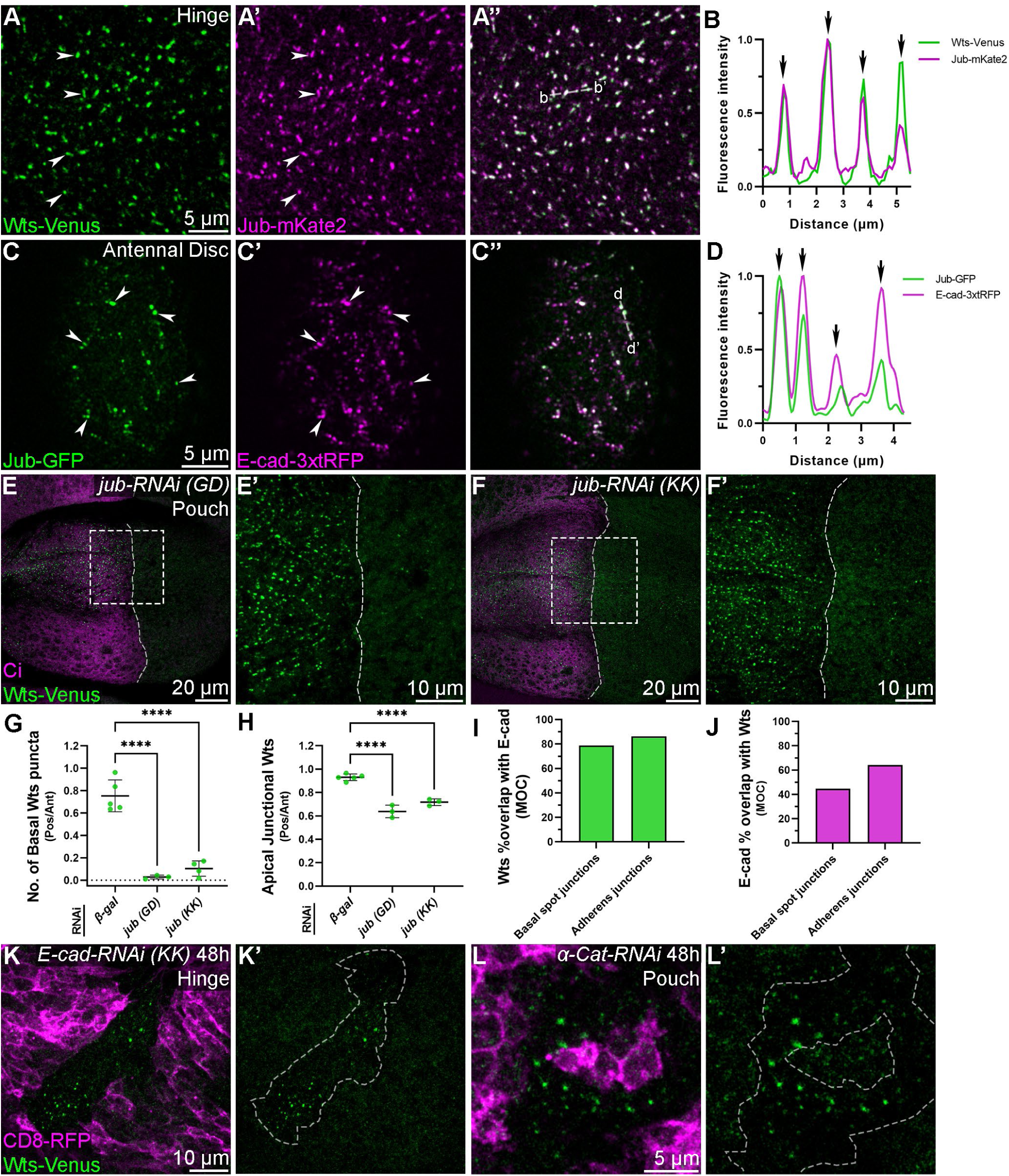
Ajuba is required for Warts to localise to basal spot junctions. **(A)** Confocal microscope images of the basal hinge region of a third instar larval wing imaginal disc. Wts-Venus is green and Jub-mKate2 is magenta. Dashed line in (**A’’**) indicates the region quantified in (**B**). Overlap between Wts-Venus and Jub-mKate2 is marked (white arrowheads in **A and A’**). **(B)** Line profile analysis of Wts-Venus and Jub-mKate2 at basal puncta. Spatial correlation of fluorescence signal is indicated (black arrows). **(C)** Super-resolution Airyscan images of the basal region of a third instar larval antennal disc. Jub-GFP is green and E-cad-3xtRFP is magenta. Dashed line in (**C’’**) indicates the region quantified in (**D**). Overlap between Wts-Venus and Jub-mKate2 is marked (white arrowheads in **C and C’**). **(D)** Line profile analysis of Jub-GFP and E-cad-3xtRFP at basal puncta. Spatial correlation of fluorescence is indicated (black arrows). **(E and F)** Confocal microscope images of the basal pouch region of a third instar larval wing imaginal disc. Wts-Venus is green and antibody-detected Ci marks the anterior compartment (magenta). The indicated jub-RNAi was expressed in the posterior compartment under the control of *en-GAL4*. Boxed regions are shown at higher magnification in (**E’** and **F’**). Dashed lines indicate the anterior-posterior compartment boundary. **(G and H)** Charts displaying the relative number of basal Wts-Venus puncta (**G**) or apical junctional Wts (**H)** present in the posterior compared to anterior pouch compartment of wing imaginal discs expressing the indicated RNAi transgenes. n = 5, 3, 4 (**G**) and 5, 3, 3 (**H**). Data are represented as mean ± SD, p values were obtained using a one-way ANOVA and Dunnett’s multiple comparison tests, **** = p<0.0001. **(I and J)** Charts showing Mander’s Overlap Co-efficient (MOC) of Wts-Venus and E-Cad-3xtRFP at basal spot junctions and adherens junctions of cells in the pouch region of third instar larval wing imaginal discs. **(K-L)** Confocal microscope images of the basal regions of the hinge (**K**) or pouch (**L**) of third instar larval wing imaginal discs. Wts-Venus is green and CD8-RFP (magenta) marks clones expressing the indicated RNAi transgenes for 48 h. Dashed lines indicate clone boundaries in (**K’ and L’**). Scale bars are indicated in image panels. All images are maximum intensity projections except for (**A-A’’**), which are single z-slices. Data is representative for 3-10 specimens per condition. See also Figure S3.

We then investigated the relationship of the basal spot junction Wts pool with key AJ proteins. Correlation studies in the wing pouch revealed that ∼80% of the basal Wts signal overlapped with E-cad, while ∼50% of the basal E-cad signal was Wts-positive and these values were similar at AJs (Figures 3I-3J). E-cad is responsible for Wts’ ability to localize to basal spot junctions, as E-cad RNAi caused both the basal spot junction and AJ Wts pools to be depleted (Figures 3K and 3K’ and S3F-S3H’). Likewise, RNAi-mediated depletion of α-catenin abolished the majority of Wts present at both basal spot junctions and AJs (Figures 3L and 3L’ and S3I and S3I’). Collectively, these experiments identify basal spot junctions as a feature of *D. melanogaster* imaginal discs and reveal a previously unidentified pool of Wts in epithelial cells in vivo. Further, they raise the possibility that Wts is recruited to at least two subcellular regions to be held in an inactive state; the AJs and the basal spot junctions.

### Warts kinase is released from basal spot junctions to be activated apically

To explore the basal spot junction pool of Wts protein further, we assessed the subcellular localisation of different Hippo pathway proteins using a number of fluorescently-tagged lines that we generated by CRISPR-Cas9 genome editing ^21–23^ or were generated using BAC transgenics ^13,14^. Consistent with previous studies ^12–14^, Hpo, Sav, Mer, Ex, Kibra and Mats, were all observed at apical junctions and/or the medial apical cortex of imaginal disc epithelial cells, where they are thought to activate Wts (Figures 4A-4D and S4A-S4G). Like Wts, each of these proteins was also observed to different degrees in the cytoplasm, with Hpo being the most cytoplasmic and Ex the least (Figure 4A-4D). However, in contrast to Wts and Jub, none of these other Hippo pathway proteins were observed at basal spot junctions (Figures 4A-4D and data not shown).

**Figure 4.**
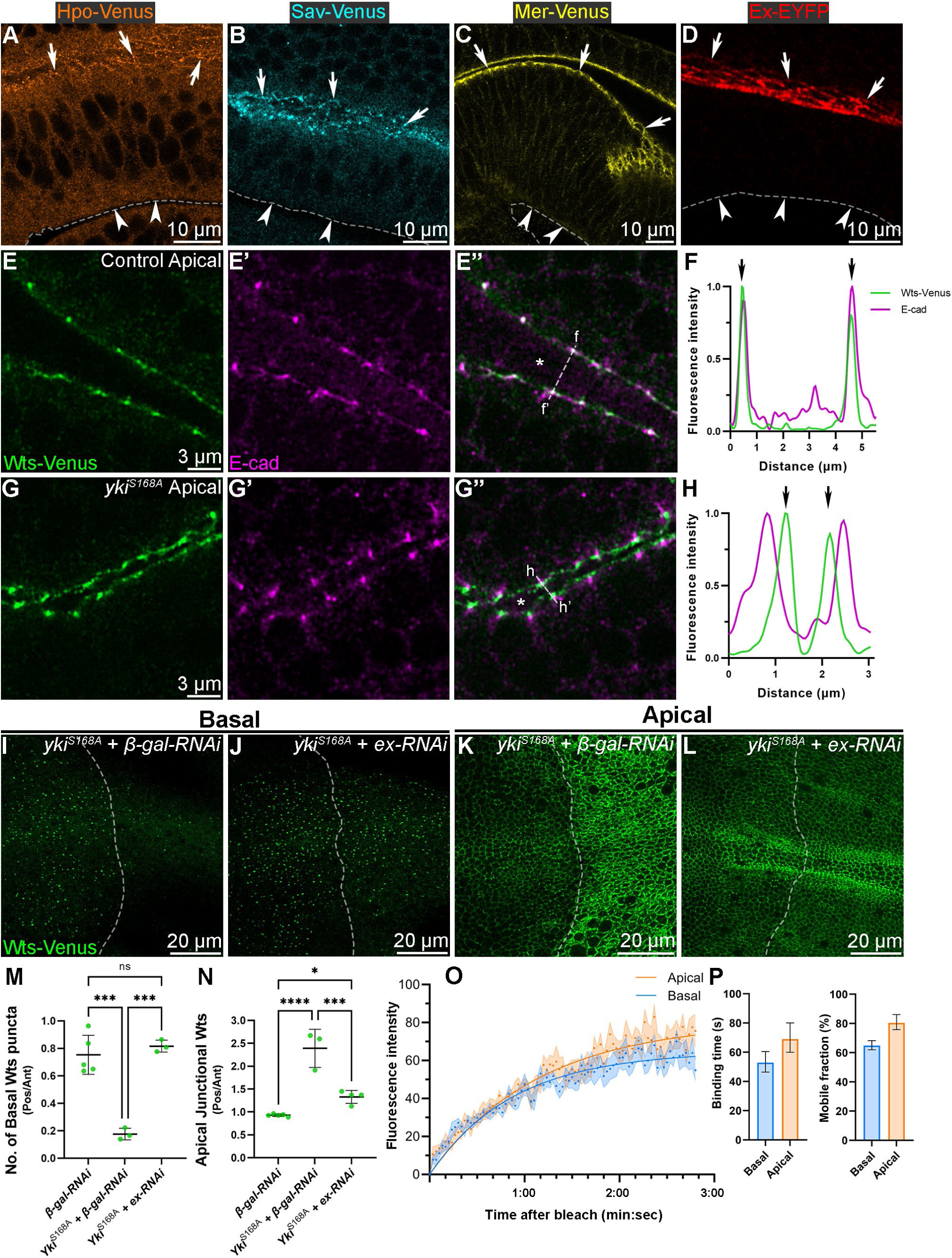
Hippo pathway hyperactivation recruits Warts away from basal spot junctions and adherens junctions. **(A-D)** Confocal microscope images showing cross-sections of third instar larval wing imaginal discs expressing either Hpo-Venus (**A**, orange), Sav-Venus (**B**, cyan), Mer-Venus (**C**, yellow) or Ex-EYFP (**D**, red). White arrows indicate the apical membrane domains and white arrowheads, and dashed lines indicate the basal membrane domains. **(E and G)** Super-resolution Airyscan images of the apical cell junctions in a fold between the hinge and pouch regions of a third instar larval wing imaginal disc. Wts-Venus is green and antibody-detected E-cad is magenta. The control anterior region of the tissue is shown in (**E**) and the posterior region, which expressed Yki^S168A^ for 24 h under the control of *en-GAL4* is shown in (**G**). The apical luminal space is marked (‘*’ in **E’’ and G’’**). Dashed lines in (**E’’ and G’’**) indicate the regions quantified in (**F and H**). **(F and H)** Line profile analyses of Wts-Venus and E-cad at AJs. Precise spatial correlation of Wts-Venus and E-cad fluorescence is marked (black arrows in **C**). Following 24 h of Yki hyperactivation, Wts-Venus is spatially separated from E-cad (black arrows in **E**). **(I-L)** Confocal microscope images of Wts-Venus (green) in either basal (**I** and **J**) or apical (**K** and **L**) regions of third instar larval wing imaginal discs. Dashed lines indicate the anterior-posterior compartment boundaries. Both Yki^S168A^ and the indicated RNAi transgenes were expressed in the posterior compartment for 24 h prior to dissection, under the control of *en-GAL4*. **(M and N)** Charts displaying the relative number of basal Wts-Venus puncta (**M**) or apical junctional Wts (**N)** present in the posterior compared to anterior pouch compartment of wing imaginal discs expressing the indicated transgenes. n = 5, 3, 3 (**M**) and 5, 3, 4 (**N**). Data are represented as mean ± SD, p values were obtained using a one-way ANOVA and Dunnett’s multiple comparison tests, ns = not significant, * = p<0.05, *** = p<0.001, **** = p<0.0001. **(O)** Chart of normalized FRAP curves of Wts-Venus in pupal notum tissues; basal is blue and apical is orange. The chart shows mean (dots) ±1 S.E.M. Solid lines indicate one-component exponential fits to data. n = 13 (basal) and 14 (apical) FRAP experiments. **(P)** Charts of FRAP-derived binding durations of Wts-Venus (left chart) or mobile fraction of Wts-Venus (right chart); basal is blue and apical is orange. Graphs show values from one-component exponential decay ±95% C.I. Scale bars are indicated in image panels. All images are single z-slices except for (**I-L**), which are maximum intensity projections. Data is representative for 3-10 specimens per condition. See also Figure S4.

In growing wing imaginal discs, Wts is not readily detectable by confocal microscopy at the sub-apical region, where it is activated by other Hippo pathway proteins^13^. Detection of endogenous Wts in this particular subcellular domain requires hyperactivation of the Hippo pathway, which can be invoked by Yki overexpression, which triggers Hippo pathway negative feedback signalling ^13^. In agreement with previous studies ^13^, expression of a transgene encoding a hyperactive Yki protein (Yki^S168A^) in the posterior wing disc for 24 h caused Wts to transition from AJs to the sub-apical region of cells (Figures 4E-4H). In addition, the basal spot junction Wts pool was substantially reduced upon expression of Yki^S168A^, indicating that this Wts pool is also sensitive to fluctuations in Hippo pathway activity in a manner analogous to AJ-associated Wts (Figures 4I and 4M). Previous studies reported that Ex is important for the Hippo feedback loop driven re-distribution of Wts from AJs to the sub-apical region ^13^. To test whether Ex depletion also impacts the basal spot junction Wts pool, we expressed Yki^S168A^ for 24 hours in wing imaginal discs together with an Ex RNAi transgene. Depletion of Ex strongly suppressed the recruitment of Wts-Venus to the apical junctions and replenished the basal spot junction pool of Wts (Figures 4J-4N). This suggests that in growing epithelial tissues, Wts can readily transit between both basal spot junctions and AJs (where it is inactive) and the sub-apical region (where it is activated).

To address this further, we employed fluorescence recovery after photobleaching (FRAP) to investigate the biophysical properties of Wts at both AJs and basal spot junctions. We performed these studies in the pupal notum as it is amenable to live imaging ^36^. First, we assessed whether basal spot junction-associated Wts was present in the pupal notum epithelia, using the clonal method described above. In the pupal notum, *wts-Venus* clones exhibited basal puncta whilst control RFP clones did not, thus validating the presence of basal spot junction-associated Wts in this tissue (Figure S5A-S5B’’’). Furthermore, basal spot junction-associated Wts co-localised with the same set of proteins as in larval wing discs, i.e., E-cad and Jub, and were present in the same focal plane as the focal adhesions (marked by Rhea), but Wts was juxtaposed to Rhea, as in wing discs (Figures S5C-S5E’’’ and S6I-6L’’). This indicates that, similar to the situation in larval imaginal discs, basal spot junctions form at the basal-most region of the lateral membranes of the pupal notum epithelia.

To perform FRAP experiments, we used a 514nm laser to bleach Wts-Venus at either basal spot junctions or regions of the AJs and then live imaged for 3 minutes to assess the degree of recovery of Wts-Venus fluorescence (Video S2). Interestingly, the recovery curves for both subcellular sites were similar: approximately 80% of the Wts-Venus signal recovered in 3 minutes at AJs, compared to 65% at basal spot junctions (Figures 4O and 4P). The binding time of Wts-Venus at AJs was calculated as 53 seconds, while at basal spot junctions it was 69 seconds (Figure 4P). Therefore, both AJ and basal spot junction pools of Wts are comprised of both immobile and mobile pools of Wts protein, where immobile refers to protein that binds for longer than 3 minutes. This indicates that the majority of Wts that resides at both AJs and basal spot junction turns over within minutes, as opposed to being stably bound at these sites. This finding is consistent with our genetic experiments that showed that Wts can be manipulated to accumulate at either the sub-apical region or basal spot junctions and AJs (Figure 4E-L). Collectively, these experiments are consistent with the possibility that Wts transits between basal and apical sites rather than being statically localized at one subcellular region.

### Warts associates with basal spot junctions of epithelial cells with enriched basal-medial actomyosin

In the course of our studies, we observed that basal Wts puncta were more prominent in specific regions of larval wing imaginal discs. In particular, they were enriched in the wing pouch, along the dorsal-ventral boundary, and also in sub-regions of the hinge epithelium, such as the dorsal anterior tissue (Figures 1G and S2B). Given that Jub and Wts are recruited to AJs in response to mechanical forces ^12,37^, we sought to determine whether basal spot junction-associated Wts correlated with regions of heightened cytoskeletal tension. We assessed basal actomyosin in wing discs using a Lifeact-tRFP transgene driven by *ubi-GAL4* and found that basal actin accumulated in the sub-regions of this tissue that also contained numerous cells with basal spot junction-associated Wts (Figures 5A-5A’’). High resolution spatial analysis of Wts and actomyosin revealed that Wts was present at basal spot junctions was distributed in between bundles of basal-medial actin (Figure 5A’’’). We then assessed the localisation of a fluorescently-tagged component of non-muscle myosin II (MyoII), an important actin-crosslinking enzymatic protein complex with contractile functions ^38^, and its relationship with basal spot junction-associated Wts. Similar to Lifeact, MyoII-3xmKate2 was enriched in the basal medial domain of wing discs and was juxtaposed to Wts at basal spot junctions (Figures 5B-5C’). Depending on the region of the wing disc imaged, the basal-medial actomyosin was arranged in different distributions; these included a donut-like organisation with radial astral filaments emanating towards basal spot junction-associated Wts (Figure 5B’), or fibre-like bundles stretched between Wts at basal spot junctions (Figure 5C’). Overall, these data are suggestive of a relationship between the accumulation of basal-medial actomyosin and the recruitment of Wts to basal spot junctions.

**Figure 5.**
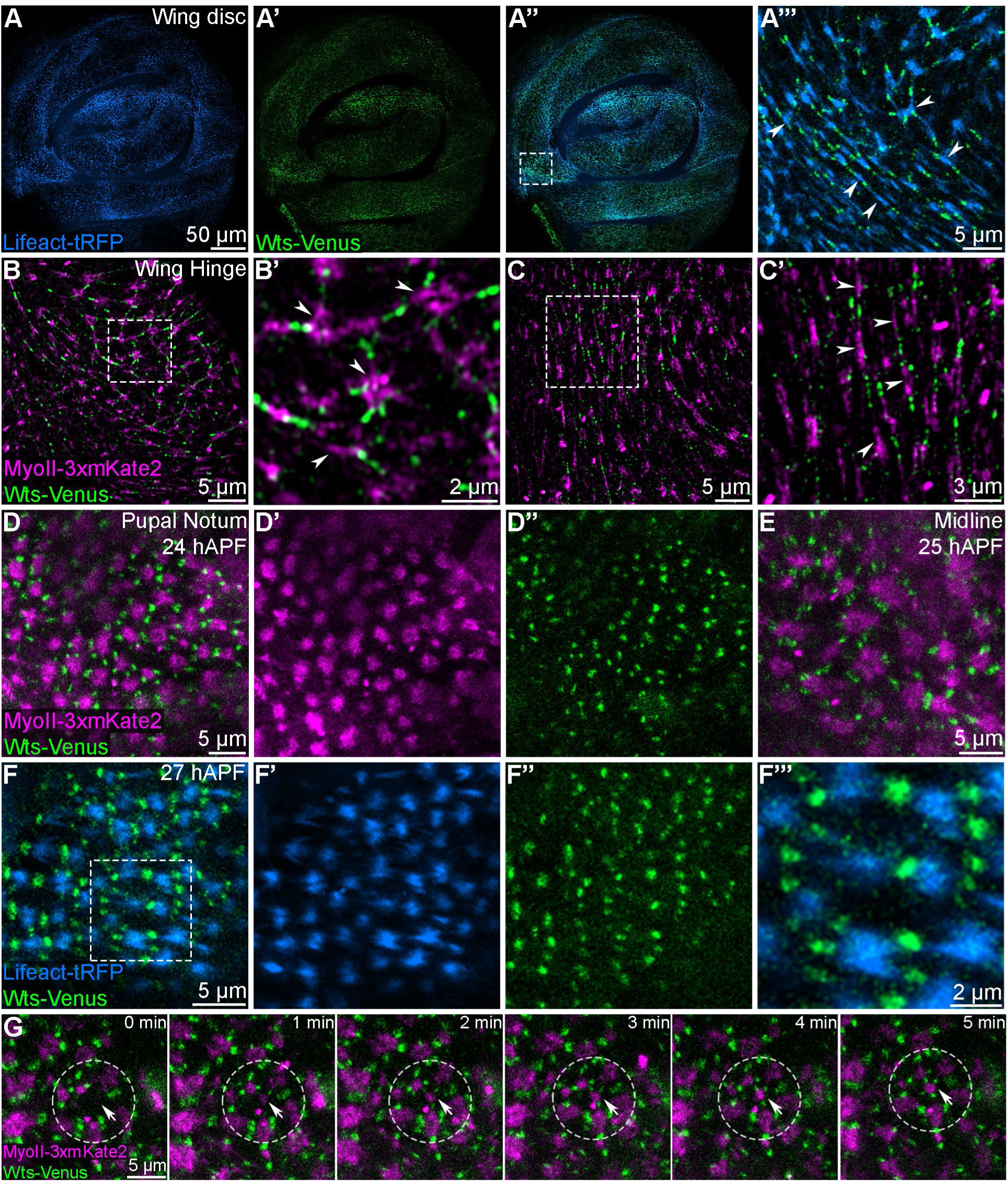
Warts protein is prominent in basal spot junctions in cells containing dense basal-medial actomyosin. **(A-C)** Confocal microscope images of the basal regions of third instar larval wing imaginal discs. Lifeact-tRFP is blue, Wts-Venus is green and MyoII-3xmKate2 is magenta. Super-resolution Airyscan images of the basal hinge regions are shown in (**B and C**). Boxed regions are displayed at higher magnification. Lifeact-tRFP is expressed ubiquitously under the control of *ubi-GAL4*. Basal spot junction-associated Wts puncta are distributed in between basal-medial actomyosin accumulations (white arrowheads in **A’’’, B’ and C’**). **(D-F)** Live confocal microscope images of the basal plane of epithelial cells in the central-posterior portion of the developing pupal notum, captured at the indicated times after puparium formation (hAPF). Wts-Venus is green, MyoII-3xmKate2 is magenta and Lifeact-tRFP is blue. Boxed region in (**F**) is shown at higher magnification (in **F’’’**). **(G)** Still images from live time-lapse confocal microscopy of the basal region of the pupal notum epithelia at 25 hAPF. A region of interest is marked (dashed circle), and the basal region of an individual cell is indicated (white arrows). The images show dynamic changes in MyoII-3xmKate2 and Wts-Venus in the basal region of the indicated cell at minute intervals. Scale bars are indicated in image panels. All images are maximum intensity projections except for (**A’’’** and **B-B’**), which are single z-slices, and (**A-A’’**), which are sum projections. Data is representative for at least 5 specimens per condition. See also Figures S5 and S6.

We hypothesized that the distribution of basal spot junction-associated Wts would change over the course of wing development, in particular during the third larval instar stage when the wing epithelia undergoes a dramatic growth phase and morphogenetic forces fold the tissue into a characteristic shape ^39,40^. To investigate this, we assessed Wts-Venus, E-cad-3xtRFP and MyoII-3xmKate2 localization in defined stages of larval wing development^41^. At the start of third instar larval development, basal spot junction-associated Wts was rare and present mostly in the presumptive pouch region of the wing disc (Figures S6A and S6A’). The abundance of basal spot junction-associated Wts puncta increased by 12 h into third instar development and continued to increase by mid to late third instar (24 h and 48 h, respectively), especially in the pouch region and in parts of the hinge (Figures S6B-S6D’). Throughout the stages of wing development studied, the distribution of basal spot junction-associated Wts correlated with the appearance of E-cad and regions of increased basal MyoII-3xmKate2, supporting the idea that basal actomyosin contractility influences recruitment of Wts to basal spot junctions (Figures S6E-S6H).

To explore the relationship between basal spot junction-associated Wts and basal actomyosin further, we utilised the pupal notum, where tissue stresses have been identified at specific stages of development ^36,37^. Using time-lapse confocal microscopy, we then characterised the dynamics of basal spot junction-associated Wts and its tissue-wide distribution during pupal notum development. In the first 18 h after pupa formation (hAPF), Wts was not readily observable, in basal spot junctions but between ∼18-26 hAPF, when cells proliferate and elongate as tensile mechanical stress increases in the posterior and central regions of the notum ^37^, a striking enrichment of basal spot junction-associated Wts was observed (Figures S61-S6K’’). Towards the end of this developmental period, particularly by ∼24 hAPF, basal spot junction-associated Wts was often neatly arranged in rows distributed along the medial-lateral axis (Figures S6K-S6K’’). Subsequently, Wts started to dissipate from basal spot junctions between 24.5 and 29.5 hAPF, whilst Rhea persisted at the basal surface of cells (Figures S6I-S6L’’).

To better characterize the dynamic relationship between basal actomyosin and basal spot junction-associated Wts, we imaged live pupal notum tissues expressing Wts-Venus and transgenic markers of actomyosin. We found that, in a manner similar to larval wing imaginal discs, basal Wts puncta were juxtaposed to rich pools of basal actomyosin in the notum (Figures 5D-5F’’’). The distribution of basal actomyosin in individual cells often had a plaque-like morphology in the basal-medial domain of each cell, surrounded by Wts puncta, but were also observed with fibrous actin protrusions stretched towards basal spot junction-associated Wts (Figures 5D-’’’). To further investigate the dynamics of basal spot junction-associated Wts and myosin in the pupal notum at 25 hAPF, and to assess whether a dynamic pulsatile actomyosin behaviour occurred in this tissue, we performed fast time-lapse confocal microscopy. However, unlike what has been reported in the egg chamber ^42^, we found that most of the basal-medial actomyosin meshwork appeared to be relatively stable once established, as did the basal spot junction-associated Wts (Video S3). We did, however, observe some interesting dynamic events, particularly whilst the basal junctions were forming and establishing their typical organisation across the tissue. For instance, we observed enrichment of an actomyosin meshwork at the basal surface of one cell, followed closely by the recruitment of multiple Wts puncta to basal spot junctions (Figure 5G). These time-lapse data show that the accumulation of basal actomyosin and Wts at basal spot junctions are highly correlated events, and can occur relatively rapidly, as we show here over a period of ∼ 5 mins (Video S4).

### Warts protein at basal spot junctions is responsive to modulation of cytoskeletal tension

Numerous studies in vertebrates and invertebrates have reported that the Hippo pathway is regulated by mechanical forces ^8,43^. For example, increased cytoskeletal tension promotes nuclear accumulation of the transcriptional co-activators YAP and TAZ (in vertebrates) and Yki (in *D. melanogaster*) and their ability to drive transcription and cell proliferation ^12,44,45^. To investigate a potential role for basal spot junction-associated Wts in mechanical regulation of organ growth, we used both genetic and chemical approaches. We first reduced cytoskeletal tension specifically in the posterior region of wing imaginal discs by RNAi-mediated depletion of Rho kinase (Rok), an upstream activator of MyoII. Three independent Rok RNAi lines caused dramatic reductions in the abundance of basal spot junction-associated Wts (Figures 6A and 6C). In agreement with previous reports ^12^, we also observed that AJ-associated Wts was reduced in these settings, but to a much lesser extent (Figures 6B and 6D). Consistent with this, RNAi-mediated depletion of a key component of MyoII, the light regulatory chain protein Spaghetti Squash (Sqh), also reduced basal spot junction-associated Wts (Figures 6E and 6E’). Next, we acutely disrupted myosin activity by bathing larval wing imaginal discs in a solution containing the Rok inhibitor, Y-27632, for 30-minutes, which caused the complete loss of basal spot junction-associated Wts, and strongly depleted basal MyoII-3xmKate2, compared to DMSO-treated control tissues (Figures 6F-6I’). Interestingly, 30-minute Rok inhibitor treatment reduced, but did not abolish, the apical junctional pool of both MyoII and Wts (Figures 6J-6K’). We also found that even a 5-minute incubation with Rok inhibitor caused a near-complete loss of basal spot junction-associated Wts, but had a milder effect on AJ-localised Wts (Figures 6F, 6G and 6J-6M). This indicates that the pool of Wts that resides in basal spot junctions of epithelial cells is highly sensitive to cytoskeletal tension and is possibly important for the ability of mechanical forces to modulate Hippo pathway activity and hence tissue growth.

**Figure 6.**
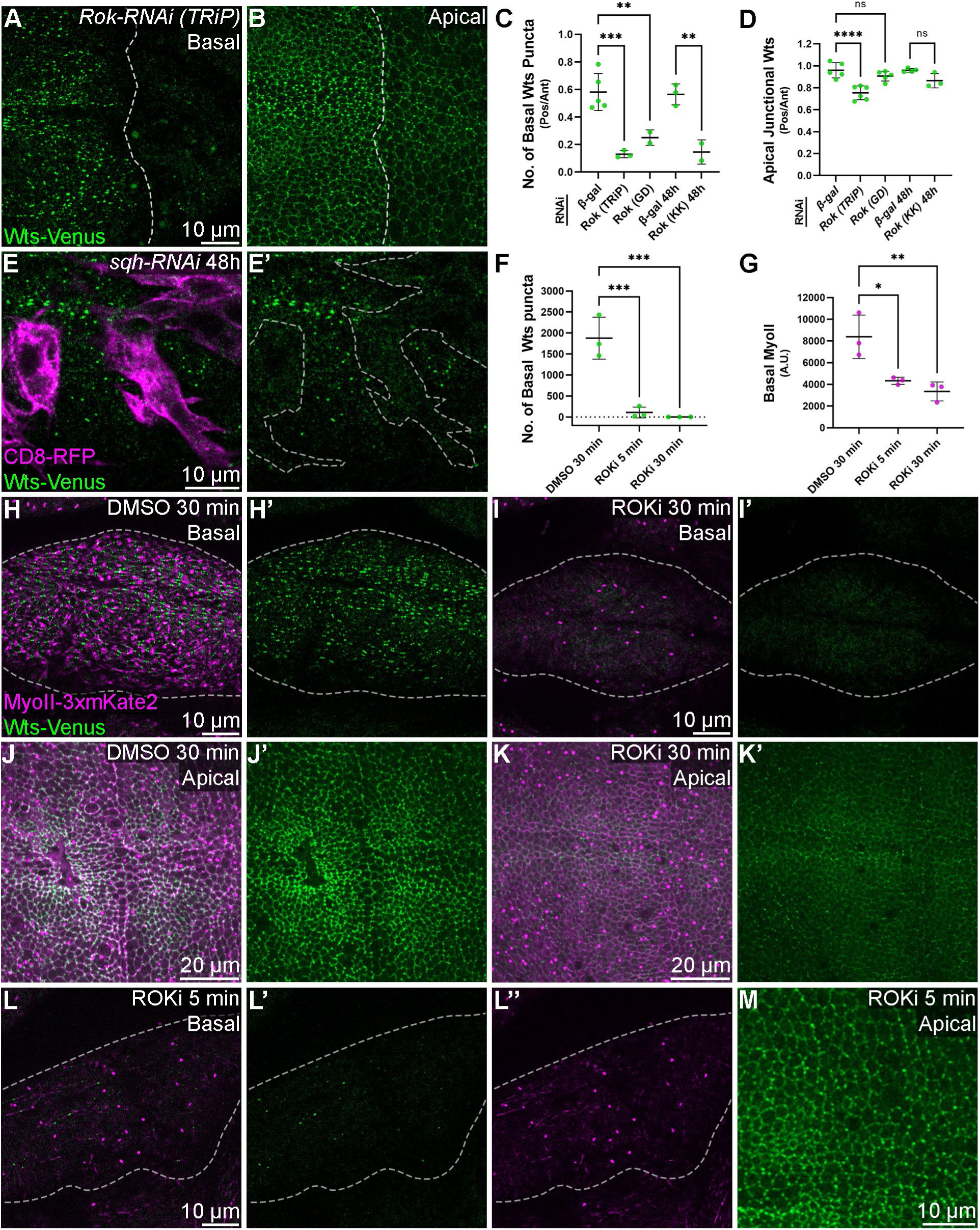
Warts protein in basal spot junctions is responsive to cytoskeletal tension. **(A-B)** Confocal microscope images of Wts-Venus (green) in the basal (**A**) or apical (**B**) pouch region of third instar larval wing imaginal discs. Rok-RNAi was expressed in the posterior compartment (right side of the dashed line, which indicates the anterior-posterior boundary of each tissue) under the control of *en-GAL4*. **(C and D)** Charts displaying the relative number of basal Wts-Venus puncta (**C**) or apical junctional Wts (**D)** present in the posterior compared to anterior compartment of wing imaginal discs expressing the indicated RNAi transgene. n = 5, 3, 2, 3, 2 (**C**) and 5, 6, 5, 4, 3 (**D**). Data are represented as mean ± SD, p values were obtained using a one-way ANOVA and Šídák’s multiple comparisons tests, ns = not significant, ** = p<0.01, *** = p<0.001, **** = p<0.0001. **(E)** Confocal microscope images of the basal pouch region of a third instar larval wing imaginal disc. Wts-Venus is green and CD8-RFP (magenta) marks clones that expressed sqh-RNAi for 48h. Dashed lines indicate clone boundaries in (**E’**). **(F and G)** Charts displaying basal Wts-Venus puncta (**F**) or basal MyoII (**G**) present in wing imaginal discs treated with either DMSO or ROKi for the indicates times. n = 3 for each condition. Data are represented as mean ± SD, p values were obtained using a one-way ANOVA and Dunnett’s multiple comparison tests, * = p<0.05, ** = p<0.01, *** = p<0.001. **(H-M)** Confocal microscope images of the pouch region of third instar larval wing imaginal discs, at the plane indicated. Wts-Venus is green and MyoII-3xmKate2 is magenta. Dashed lines in (**H-I’ and L-L’’**) indicate the basal pouch region. Tissues shown were bathed in media containing either DMSO (**H and J**) or 1 mM Rok inhibitor, Y-27632 (**I, K, L and M**) for the durations indicated prior to dissection. Scale bars are indicated in image panels. All images are maximum intensity projections except for (**E-E’**, **H-I’**, and **L-L’’**), which are single z-slices. Data is representative of at least 2-6 specimens per condition. See also Figures S5 and S6.

### Focal adhesions influence Warts recruitment to basal spot junctions

Focal adhesions are macromolecular assemblies that link cells to the ECM via integrin receptors and connect to the intracellular cytoskeletal network via actin binding proteins such as Rhea ^4,5^. Given the close proximity of basal spot junction-associated Wts and basal focal adhesions, we further investigated potential relationships between them. In both larval wing imaginal discs and in the pupal notum, we found that the basal-medial actin network overlapped with focal adhesions (Figure 5A’’’ and Figures S5F-S5G). These observations suggest that basal-medial actomyosin assemblages are physically linked to both focal adhesions and basal spot junctions. Next, we determined whether perturbation of focal adhesions impacted basal spot junction-associated Wts, by depleting the key focal adhesion protein Rhea, which is required to link integrins to the cytoskeleton, and is essential for focal adhesion formation^46^. Rhea RNAi was expressed using *en-GAL4* and GAL80ts for 48 h immediately prior to tissues dissection as longer expression caused organism lethality. Transient Rhea depletion in the posterior region of the larval wing imaginal disc (Figure S7A-B and E), caused a striking 10-20-fold increase in the association of Wts at basal spot junctions (Figures 7A, 7E and S7G-H’). This was most prominent in the ventral and dorsal hinge regions of the wing disc – sites that in control tissues normally display few, if any, basal Wts puncta (see Figures 1G and S2B). The altered distribution of Wts in these tissue regions coincided with an increase in the size and intensity of basal spot junctions, as assessed by basal E-cad-3xtRFP (Figure 7A’), demonstrating that basal spot junctions can be induced when cell-ECM attachment is perturbed. Consistent with this, transient RNAi-mediated depletion of either of the two *D. melanogaster* integrin subunits, Myospheroid (Mys) and Inflated (If), caused a striking increase in Wts and E-cad at basal spot junctions (Figures 7B, 7B’, 7E, S7C-D, S7F and S7I-J’). By contrast, transient depletion of either Rhea or Mys caused a minor reduction in AJ-associated Wts (Figure S8A-S8F), and the localisation of other hippo pathway proteins, such as Hpo-Venus, were not obviously impacted by transient Rhea depletion (Figure S7M-N’).

**Figure 7.**
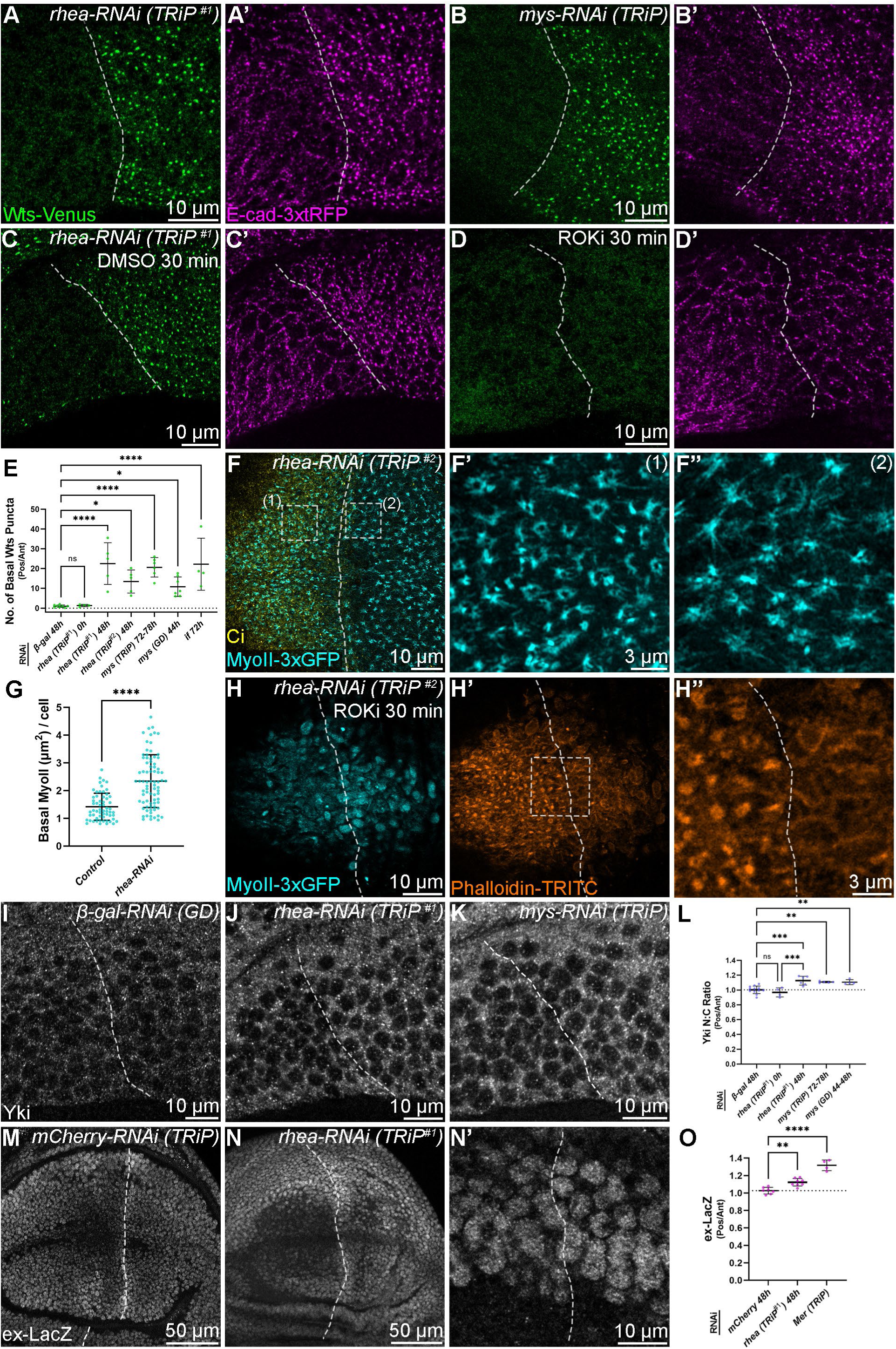
Focal adhesions influence Warts recruitment to basal spot junctions and Hippo pathway-mediated repression of Yorkie. **(A-D)** Confocal microscope images of the basal surface of the ventral pouch-hinge boundary fold in third instar larval wing imaginal discs. Wts-Venus is green, E-cad-3xtRFP is magenta. rhea-RNAi (**A** and **C-D**) or mys-RNAi (**B**) transgenes were expressed for 48 h in the posterior compartment under the control of *en-GAL4*. In (**C-D**), the wing imaginal discs were bathed for 30 min in media containing either DMSO or 1 mM Rok inhibitor, Y-27632, prior to imaging. Dashed lines indicate the anterior-posterior boundaries, posterior is to the right. **(E)** Chart displaying the relative number of basal Wts-Venus puncta present in the posterior compared to anterior compartment of the ventral pouch-hinge boundary epithelia in wing imaginal discs expressing the indicated RNAi lines for the indicated times. n = 8, 4, 5, 5, 6, 7, 4. Data are represented as mean ± SD, p values were obtained using a one-way ANOVA and Dunnett’s multiple comparison tests, ns = not significant, * = p<0.05, **** = p<0.0001. **(F)** Super-resolution Airyscan images of the basal pouch region of a third instar larval wing imaginal disc. MyoII-3xGFP is cyan and antibody-detected Ci marks the anterior compartment (yellow). Dashed lines indicate the anterior-posterior boundary. rhea-RNAi was expressed for 48 h in the posterior compartment under the control of *hh-GAL4*. Boxed regions are shown at higher magnification. **(G)** Chart showing quantification of the basal-medial cell area, marked by MyoII-3xGFP, in cells expressing Rhea-RNAi compared to control cells. Two-tailed Mann-Whitney test; n=59, 79; error bars indicate mean and standard deviation, ****P≤0.0001. **(H)** Confocal microscope images of the basal pouch region of a third instar larval wing imaginal disc. MyoII-3xGFP is cyan, Hoechst marks DNA (blue), antibody-detected Ci is yellow and Phalloidin-TRITC is orange. Dashed lines indicate the anterior-posterior boundary. rhea-RNAi was expressed for 48 h in the posterior compartment under the control of *hh-GAL4* and subsequently bathed for 30 min in media with 1 mM Rok inhibitor, Y-27632. Boxed region is shown at higher magnification. **(I-K)** Confocal microscope images of third instar larval wing imaginal discs, antibody-detected Yki is greyscale. The indicated RNAi lines were expressed for 48 h in the posterior compartment under the control of *en-GAL4*. Dashed lines indicate the anterior-posterior boundary, posterior is to the right. **(L)** Chart showing the nuclear-cytoplasmic ratio of Yki in posterior compared to anterior wing discs expressing the indicated RNAi lines. n=12, 4, 6, 5, 4. Data are represented as mean ± SD, p values were obtained using a one-way ANOVA and Dunnett’s multiple comparison tests, ** = p<0.01, *** = p<0.001. (**M-N’**) Confocal microscope images of third instar larval wing imaginal discs, transcriptional activity of the *ex* gene was detected by β-gal antibodies (greyscale). The indicated RNAi lines were expressed for 48 h in the posterior compartment under the control of *hh-GAL4*. Dashed lines indicate the anterior-posterior boundary, posterior is to the right. **(O)** Chart comparing the ratio of β-gal in posterior compared to anterior wing discs expressing the indicated RNAi lines. n=7, 8, 4. Data are represented as mean ± SD, p values were obtained using a one-way ANOVA and Dunnett’s multiple comparison tests, ** = p<0.01, **** = p<0.0001. Scale bars are indicated in image panels. All images are maximum intensity projections except for (**I-K** and **M-N’**), which are single z-slices. Data is representative of at least 4 specimens per condition. See also Figures S7, S8 and S9.

Given our observation that actomyosin contractility is critical for the recruitment of Wts-Venus to basal spot junctions, we hypothesised that altered myosin activity underlies the enhanced recruitment of Wts to basal spot junctions in Rhea-depleted cells. To test this, we bathed Rhea-depleted wing discs in solution containing the Rok inhibitor, Y-27632, for 30 minutes and found that Wts was completely lost from basal spot junctions (Figures 7C and 7D). Furthermore, the increase in basal spot junctions themselves following Rhea depletion was also reversed, as determined by assessing basal E-cad (Figures 7C’ and 7D’). This indicates that in cells with compromised focal adhesions, actomyosin contractility at the basal-medial cell surface stabilizes basal spot junctions and causes increased recruitment of Wts to them. To investigate this further, we assessed the distribution of the basal-medial actomyosin network in these cells by imaging fluorescently-tagged Sqh (MyoII-3xGFP) and found that the basal myosin organisation was dramatically altered. Compared to control wing pouch epithelial cells, where myosin formed a characteristic donut-like organisation with filamentous projections emanating radially, in Rhea-depleted cells, myosin was organised in an astral formation and was distributed across a larger basal-medial cell surface area (Figures 7F-7G). These results indicate that focal adhesions have a profound influence on the organisation of the basal actomyosin network, similar to observations reported in both the *D. melanogaster* follicular epithelium ^47^ and embryo^48^. Strikingly, when Rhea was depleted by RNAi and cells were treated with Rok inhibitor, F-actin was largely depleted from the basal medial domain and was mostly cortical (Figures 7H-7H’’). By comparison, control cells with unperturbed focal adhesion complexes retained a strong pool of F-actin at the focal adhesion sites (Figure 7H-7H’’). Furthermore, the disorganised, bulging morphology of Rhea-depleted and Rok inhibited tissues suggests that both basal actomyosin and cell-ECM attachment via focal adhesions are critical for cell morphology at the basal part of the wing disc epithelium.

Finally, we considered whether increased recruitment of Wts to basal spot junctions would impact Hippo pathway activity. To investigate this, we employed the Grab-FP technique ^49^, which allows GFP-tagged proteins to be re-localized to either basolateral or apical regions of the cell. Forced re-localization of the majority of Wts-Venus to basolateral cell membranes caused mild wing overgrowth, while apical re-localisation of Wts-Venus moderately suppressed wing growth (Figures S9A-S9G). Consistent with this, basolateral re-localization of Wts-Venus increased expression of the Yki reporter *ex-lacZ*, whilst apical re-localization of Wts-Venus decreased *ex-lacZ* (Figures S9H-S9K). This is consistent with our previous data that suggested that basal spot junctions are a site of Wts repression. However, the limitations with these experiments are that basolateral Grab-FP re-localized Wts-Venus to a broader basolateral membrane domain than just basal spot junctions, and actually reduced basal-spot junction associated Wts. Similarly, a caveat with the apical Grab-FP experiment is that it could theoretically reduce both the AJ and basal spot-junction pools of Wts-Venus, not just the basal pool.

Therefore, to explore a role for basal spot junctions in Wts repression further, we took advantage of the striking increase in basal spot junction-associated Wts in Rhea-depleted cells. If, as predicted, Wts activity is repressed when localised at basal spot junctions, transient Rhea depletion might cause an increase in Yki activity and nuclear localization, which is normally limited by Wts ^15–17^. To assess this, we immunostained larval wing discs that expressed RNAi lines for between Rhea or Mys for 44 h and 78 h with a Yki antibody and quantified the nuclear and cytoplasmic Yki signal. We assessed multiple wing discs under these conditions and observed a small but significant increase in nuclear Yki in both Rhea- and Mys-depleted cells (Figures 7I-7L and S7K). In accordance with this, *ex-LacZ* was modestly elevated upon transient Rhea protein, relative to control imaginal discs (Figures 7M-7O). Constitutive expression of a Merlin (Mer) RNAi transgene served as a positive control for this experiment and also increased *ex-lacZ* levels (Figures 7O and S7L). The observed subtle increase in Yki activity in these experiments probably reflected the fact that only a fraction of the total Wts pool was re-localised to basal spot junctions and repressed upon transient Rhea or Mys depletion and that Wts abundance is also increased in these cells (Figure S8G-S8H). These results are consistent with the hypothesis that Wts, when localised to basal spot junctions, is inactive, as it is at AJs.

## DISCUSSION

Recruitment of key proteins to specific subcellular domains is an important feature of cell signaling and is manipulated to stimulate activity changes. In *D. melanogaster* epithelial tissues, upstream Hippo pathway proteins (e.g., Ex, Mer and Kibra) and core kinase proteins (e.g., Hpo and Wts) form activating complexes at the sub-apical region and the medial apical cortex ^13,14^. By contrast, Wts associates with Jub in an inactive state at AJs ^12^. Wts localization at AJs is sensitive to cytoskeletal tension, because of tension-dependent recruitment of Jub to AJs via the mechanotransducer protein α-catenin. In this way mechanical forces are thought to control Yki activity, cell proliferation and organ size ^12,50,51^. Here, using electron microscopy, we discovered a previously undescribed pool of Wts protein at the basal-most region of lateral membranes of epithelial cells, which we term basal spot junctions. These resembled many features of apical AJs and were evident in periods of organogenesis when morphogenetic forces were prominent. Wts is likely held in an inactive complex in basal spot junctions, given that it requires Jub to stably associate in them, as it does at AJs, while other Hippo pathway proteins that activate Wts do not localize to basal spot junctions. Thus, we have identified a new subcellular region that has the potential to integrate morphogenetic forces and Hippo signalling activity. Our study further highlights the importance of control of Wts subcellular localization for Hippo pathway activity, and the competing influences of cytoskeletal tension and upstream Hippo pathway proteins on Wts localization and hence activity.

Basal spot junction-associated Wts is highly sensitive to changes in actomyosin contractility, a known modulator of Wts recruitment to AJs and Wts activity ^12^. In fact, depletion of Jub by RNAi, as well as modulation of cytoskeletal tension (either genetically or chemically), more profoundly impacted Wts localisation at basal spot junctions than at AJs. This suggests that the basal spot junction pool of Wts is more mechanosensitive than the AJ pool, or that apical AJs are more mature and stable structures, and so they are less easily disrupted by experimental perturbations of actomyosin. Alternatively, Wts might be recruited to AJs by multiple mechanisms, some of which are insensitive to mechanical forces. The localization of Wts to basal spot junctions was clearly distinct from focal adhesions, but we found that actomyosin is intimately associated with both structures. Basal actomyosin in imaginal discs was predominantly observed in one of two organizations; as bands connecting two distinct cell junctions, or in radial asters that emanated from focal adhesions and often associated with multiple sites on cell membranes. Disruption of focal adhesions caused basal actomyosin to redistribute and cover a greater area of the basal-medial cell surface and, in turn, this dramatically increased the presence of basal spot junctions and Wts at these locations. This suggests that loss of focal adhesions might cause a redistribution of mechanical forces that stabilise spot junctions. In support of this notion, we found that actomyosin contractility was essential for the increase in spot junctions observed upon focal adhesion loss. Thus, basal spot junctions appear to be stabilized by actomyosin contractility, akin to the important role that actomyosin contractility plays in AJ stabilisation ^52^.

Interestingly, in many ways, the basal spot junctions we describe here resemble features of AJs: 1) basal spot junctions are E-cad-rich and form contacts with E-cad on apposing cellular membranes; 2) they are also comprised of α-catenin and Jub; and 3) they are physically coupled to the cytoskeleton, and therefore link cell-cell adhesion and intracellular cytoskeletal tension. To our knowledge, basal spot junctions have not been reported previously in *D. melanogaster* imaginal discs, but similar structures have been reported in the *D. melanogaster* embryo. Spot adherens junctions, as opposed to belt-like continuous zonula adherens, have been observed on the lateral membranes of blastoderm, ectoderm and mesoderm cells in the early *D. melanogaster* embryo ^53,54^. Spot-like AJ’s have also been observed along the lateral membranes of cultured human epithelial cells, though cells imaged in these studies lacked the complex 3D organization of an epithelial tissue ^55–60^. As such, it will be interesting to determine whether basal spot junctions are present in different mammalian epithelia and if they are mechanoresponsive and regulate the subcellular localization of the mammalian Wts orthologues (LATS1 and LATS2) and thus Hippo pathway activity. The presence of Wts in basal spot junctions was not observed broadly in epithelial tissues throughout their development; instead Wts was present at basal spot junctions in temporally- and spatially-restricted fashions, such as the pouch and hinge regions of the late larval wing imaginal discs, and in the central-posterior region of the pupal notum 18-26h APF, when it experiences morphogenetic stresses ^37^. As such, for any investigation of basal spot junctions and Hippo signaling in mammals *in vivo*, it will be important to investigate tissues that are similarly experiencing morphogenetic stresses.

Wts represses the key transcriptional co-activator Yki, which regulates a gene expression program that drives cell proliferation and tissue growth. As such, given Hippo’s important growth-regulatory role one might predict that Wts activity would increase at the cessation of organ growth to repress Yki’s growth-promoting potential. In line with this, the nuclear abundance of Yorkie has been reported to decrease as wings grow ^61,62^. However, we observed no obvious decrease in Wts at basal spot junctions as larval wing imaginal discs grew. Rather, basal spot junction-associated Wts was largely absent in the early phases of larval wing disc growth, and more prevalent as tissues developed and underwent morphogenic events such as folding. Thus, localization of Wts to basal spot junctions cannot obviously be linked to termination of organ growth once the appropriate size has been attained. Instead, we hypothesize that cells that experience morphogenetic stresses act via basal spot junctions to repress Hippo pathway activity and in turn elevate Yki activity. Elevated Yki-regulated transcription could potentially provide a survival advantage to these cells or even stimulate their proliferation. Indeed, we observed a striking accumulation of Wts at basal spot junctions in the central-posterior region of the pupal notum 18-26h APF, which coincides with a period of cell proliferation and elongation as tensile mechanical stress increases along the medial-lateral axis and the tissue becomes anisotropic ^36,37^. Apical stress fibres in these pupal notum cells have been shown to cluster Wts at AJs and thereby reduce its activity and allow Yki-induced proliferation to occur ^37^. Our study suggests that basal spot junctions also regulate Wts and hence Yki activity in these same cells in order to influence cell proliferation in response to morphogenetic forces. Finally, the UAS-APEX2-GBP transgenic *D. melanogaster* strains generated here should allow high resolution analysis of any GFP-tagged protein that is expressed in somatic *D. melanogaster* tissues by electron microscopy.

## MATERIALS AND METHODS

### Drosophila melanogaster genetics

The following *D. melanogaster* stocks were used, some of which were from the Bloomington *Drosophila* Stock Centre (BDSC) or the Vienna *Drosophila* Resource Centre (VDRC): *wts-Venus, FRT82B wts-Venus, hpo-Venus, FRT42D hpo-Venus*, *FRT82B Kibra-Venus* and *sav-Venus* ^21^*, Mer-Venus* and *Kibra-Venus* ^22^*, GFP-wts* ^12^*, GFP-mats* ^13^*, jub-GFP* ^63^*, mTFP1-wts-mCitrine* ^27^*, ex-eYFP* ^14^, *jub-mKate2* ^37^*, sqh-3xmKate2, sqh-3xGFP and E-cad-3xtRFP* ^64^*, E-cad-GFP* ^65^*, rhea-mCherry* (BDSC, #39648)*, rhea-EGFP* (BDSC, #39649)*, α-Cat-EGFP* (BDSC, #59405), UAS-*Lifeact-tRFP* (BDSC, #58714)*, UAS-jub RNAi^GD^* (VDRC, #38443), *UAS-jub RNAi^KK^*(VDRC, #101993), *UAS-rhea RNAi^TRiP#^*^1^ (BDSC, #33913), *UAS-rhea RNAi^TRiP#2^* (BDSC, #32999), *UAS-Rok RNAi^TRiP^* (BDSC, #28797), *UAS-Rok RNAi^GD^* (VDRC, #3793), *UAS-Rok RNAi^KK^* (VDRC, #104675) *UAS-sqh RNAi^GD^* (VDRC, #7916), *UAS-α-Cat RNAi^KK^* (VDRC, # 107298), *UAS-E-cad RNAi^KK^* (VDRC, #103962), *UAS-ex RNAi^KK^*(VDRC, #109281)*, UAS-β-gal RNAi^GD^ (VDRC, #51446), UAS-mys RNAi^TRiP^ (BDSC, #33462), UAS-mys RNAi^GD^ (VDRC, #29619), UAS-if RNAi^TRiP^ (BDSC, #27544), UAS-Mer RNAi^TRiP^ (BDSC, #28007), UAS-Yki-S168A-V5* (BDSC, #28818)^66^, *UAS-myc-wts* (BDSC, #56809), *ex-lacZ* (BDSC, #44248), *en-GAL4* (BDSC, #30564), *hh-GAL4*, *tub-GAL80ts* (BDSC, #7108), *Ubi-GAL4* (BDSC, #32551), *hsFLP* (BDSC, #7), *Act5C>CD2>GAL4* (BDSC, #4780), *FRT82B ubi-mRFP-NLS* (BDSC, #30555), *UAS-mRFP-H2B*, *UAS-CD8-RFP* (BDSC, #27391), *UAS-P35* (BDSC, #5073), basolateral GrabFP (BDSC, #68175), apical GrabFP (BDSC, #68178), extracellular GrabFP (BDSC, #68174) and *UAS-APEX2-GBP* (this study). *FRT42D E-cad-GFP* and *FRT42D E-cad-3xtRFP* were generated by meiotic recombination.

*D. melanogaster* were raised at 25°C on a standard diet made with semolina, glucose (dextrose), raw sugar, yeast, potassium tartrate, nipagen (tegosept), agar, propionic acid and calcium chloride. Animals were fed in excess food availability to ensure that nutritional availability was not limiting. In some instances, animals were raised at 18°C to restrict transgene expression during early development using *tub-GAL80ts* and were subsequently transferred to 29°C to induce transgene expression.

### Generation of UAS-APEX2-GBP D. melanogaster

The APEX2-GBP cDNA was amplified by PCR from p3E-GBP ^24^ and cloned into pUAST. Transgenic *D. melanogaster* were generated by Bestgene Inc.

### Electron microscopy

*D. melanogaster* third instar larvae were dissected in ice-cold PBS; the posterior portion of the animal was removed, and the anterior portion inverted to expose the imaginal discs to solutions. Tissues were fixed for 1 hr in 2.5% glutaraldehyde in 0.1M sodium cocadylate, washed three times for 5 mins in 0.1M sodium cocadylate buffer, and then incubated with 1 mg/mL DAB in 0.1M sodium cocadylate for 15 mins. Samples were then incubated in fresh DAB solution, in the presence of 0.02% H_2_O_2_, for 30 mins to generate the insoluble APEX2 reaction product. Subsequently, tissues were washed three times for 5 mins in 0.1M sodium cocadylate, and then post-fixed in 1% OsO_4_ in 0.1M sodium cocadylate for 2 hrs at 4°C to convert the DAB reaction product into an electron-dense product surrounding the bound protein of interest. Subsequently, tissues were washed three times for 5 mins in 0.1M sodium cocadylate to remove the OsO_4_, and then washed three times for 5 mins in Milli-Q H_2_O to remove the codaylate buffer. Individual wing imaginal discs were then carefully dissected off the inverted anterior portion of the larvae, and serially dehydrated in ethanol (30%, 50%, 70%, 90%, then twice in 100%), and then transferred to 100% propylene oxide. For each of these steps, tissues were incubated three times for 1 min in a Biowave microwave at 150W. Tissues were then serially infiltrated with increasing concentrations of Epon 812 resin (25%, 50%, 75%, then three times in 100%). At each step, the samples were incubated for 3 mins in a Biowave microwave at 250W with the vacuum on. The resin was then polymerised at 65°C for 48 hrs. Ultrathin sections were cut on an ultramicrotome (Leica EM UC7, Leica Microsystems), mounted on formvar coated slot grids (ProSciTech), and imaged using a JEOL1400 electron microscope (JEOL) at 80 kV., the samples were incubated for 3 mins in a Biowave microwave at 250W with the vacuum on. The resin was then polymerised at 65°C for 48 hrs. Ultrathin sections were cut on an ultramicrotome (Leica EM UC7, Leica Microsystems), mounted on formvar coated slot grids (ProSciTech), and imaged using a JEOL1400 electron microscope (JEOL) at 80 kV.

### Immunostaining and confocal microscopy

*D. melanogaster* larval imaginal discs were dissected in PBS and fixed in 4% paraformaldehyde and either mounted directly in VectaShield Mounting Medium (Vector Laboratories, H-1000) or first permeabilised with PBST (PBS with 0.3% Triton X-100) and immunostained. The following primary antibodies were used: rat anti-Yki (1:400, Iswar Hariharan), rat anti-Ci (1:20, Developmental Studies Hybridoma Bank), rat anti-E-cad (1:20, Developmental Studies Hybridoma Bank), mouse anti-β-Gal (1:100, Sigma-Aldrich, G4644), mouse anti-Mys (1:100, Developmental Studies Hybridoma Bank, CF.6G11), and mouse anti-Arm (1:50, Developmental Studies Hybridoma Bank, N2 7A1). Secondary antibodies conjugated to Alexa Fluor 405, Alexa Fluor 568 or Alexa Fluor 647 (Invitrogen, Life Technologies, by Thermo Fisher Scientific) were used at a concentration of 1:500. Hoechst 33342 (1 ug/ml, Sigma-Aldrich, B2261) was used to stain nuclei and Phalloidin-TRITC (1:1000, Sigma-Aldrich, P1951) to stain F-actin. Images were acquired on a Zeiss LSM980 Airyscan2 microscope equipped with a 40X 1.3 NA oil immersion objective (for fixed samples) or a 40X 1.1 NA water immersion objective (for live samples). The Airyscan2 detector was used for super-resolution image acquisitions. The spectral GaAsP detector with two flanking PMT’s were used for all other confocal images. Raw Airyscan data were processed using the default parameters in ZEN software (Blue edition, Zeiss). For super-resolution data, chromatic shifts in x, y, and z dimensions were corrected using Chromagnon (v85) ^67^, using a biological calibration sample. In some instances, larval imaginal discs were bathed in 1 mM ROK inhibitor at 25°C (Y-27632, Cell Signalling Technologies, 72304) or DMSO (Sigma-Aldrich, D2650), in Schneider’s Insect Medium (Sigma-Aldrich, S0146) with 10% FBS (Sigma-Aldrich, F9423) prior to fixation, mounting and imaging.

### Live confocal microscopy

*D. melanogaster* pupae were mounted and imaged as in ^68^. Pupae were collected at puparium formation and transferred into small Petri dishes until the dissections of the nota were processed at 16–17 h after puparium formation (hAPF). In some cases, pre-pupae yet to undergo head eversion were staged at head eversion, estimated to occur at 12 hAPF. The anterior two thirds of the pupal case was removed, and the pupa was mounted dorsal side down on a drop of Halocarbon oil 700 on a no 1.5 high precision coverslip mounted in an Attofluor imaging chamber (Thermo Fisher Scientific). Pupae were stabilised with a drop of vacuum grease on the posterior tip, and the notum was imaged live using a Zeiss LSM980 Airyscan 2 microscope.

### Image analysis and statistics

Fluorescence microscopy images were analysed using ImageJ/Fiji ^69^. To quantify Yki and *ex-LacZ* (β-Gal) fluorescence, segmentation of nuclei area and cytoplasmic area was performed on representative single z-planes consisting of both control and experimental tissues by using Hoechst fluorescence to specify the nuclei area. The subcellular levels of Wts at apical and basal sites was determined by tracing individual clones containing 2 or more cells (using CD8-mRFP), and fluorescence intensity measurements above the cytoplasmic signal recorded. The number of basal Wts-Venus puncta was determined from segmented z-stack images by counting particles >1 μm in size in equal-sized regions of control and experimental tissues that together covered the entire pouch region or the ventral pouch-hinge boundary fold epithelium. Apical junctional Wts-Venus fluorescence was determined from segmented maximum intensity projection images of the pouch region, and equal-sized regions of interest (covering >200 cells) in control and experimental tissue were compared. Co-localisation analysis of E-cad-3xtRFP and Wts-Venus was performed using the JaCoP plugin ^70^. Airyscan super-resolution Z-stack data were manually thresholded to specify the regions of interest, and Manders’ overlap coefficient calculated. MyoII-3xGFP fluorescence was quantified from Airyscan super-resolution data in control and experimental tissues, at optimal focal planes, by using consistent thresholding intensity values to segment MyoII area. Total Wts protein was determined from sum projection z-stack data that covered the entire wing pouch region in 3D, and equal-sized regions of interest in the control and experimental portions of the tissue were compared. Adult wing size data was analysed using Photoshop’s magnetic lasso tool to define the anterior and posterior compartments. Line profile fluorescence intensity data was normalised, and all data were collated using Excel (Microsoft). GraphPad Prism 9 was used to generate graphs and to perform statistical analyses.

## ACKNOWLEDGEMENTS

We thank members of the Harvey lab, Alpha Yap and Ivar Noordstra for discussions and comments on the manuscript. We thank Markus Affolter, Yohanns Bellaiche, Rick Fehon, Iswar Hariharan, Ken Irvine, Gary Struhl, the Bloomington *Drosophila* Stock Center, the Vienna *Drosophila* RNAi Center, the Australian *Drosophila* Research Support Facility (www.ozdros.com), and the Developmental Studies Hybridoma Bank for *D. melanogaster* stocks and antibodies, Thomas Hall, Rob Parton and Xiaomeng Zhang for plasmids. K.F.H was supported by a Senior Research Fellowship (APP1078220) and Investigator grant (APP1194467) from the National Health and Medical Research Council of Australia. This research was supported by the Australian Research Council (DP180102044, DP190101743, DP22010523 and DP230101406). We acknowledge the Monash Micro Imaging Facility and the Monash Ramaciotti Centre for Cryo-Electron Microscopy.

