## Supplemental Data for "Basal spot junctions of epithelial tissues respond to morphogenetic forces and regulate Hippo signaling"

**Kroeger et al., Supplemental Data**

**Figure S1. APEX2-GBP overexpression does not alter the subcellular localisation of Warts-Venus.**

**(A)** Schematic representation of the APEX2-GBP system. GFP-tagged proteins of interest (POI) are bound by APEX2-GBP. The enzymatic activity of APEX2 converts DAB into an electron dense precipitate that can be visualised by electron microscopy.

**(B)** Stereo microscope image of a *D. melanogaster* third instar larval wing imaginal disc expressing APEX2-GBP in the posterior compartment under the control of *en-GAL4*. Dashed line indicates the anterior-posterior compartment boundary. APEX2 converts DAB into a high contrast, electron dense reaction product.

**(C)** Confocal microscope images of apical regions of third instar larval wing imaginal discs expressing APEX2-GBP under the control of *en-GAL4*. Wts-Venus is green and antibody-detected Ci marks the anterior compartment (magenta). Boxed region in is shown at higher magnification. Dashed lines indicate the anterior-posterior compartment boundary.

**(D-F)** Adult female *D. melanogaster* wings expressing APEX2-GBP under the control of *en-GAL4*, either in control animals (**D**) or in *wts-Venus* homozygotes (**E**). As a positive control for this experiment, a transgene encoding Myc-tagged Wts protein was also expressed (**F**) (note, these animals were reared at 18°C to avoid lethality). Anterior and posterior regions indicated in (**D**), are quantified in **(G).**

**(E)** Chart showing the ratio of posterior wing area compared to anterior wing area for the indicated genotypes. n=10 for each condition. Data are represented as mean ± SD, p values were obtained using a one-way ANOVA and Dunnett's multiple comparison tests, ** = p<0.01, **** = p<0.0001.

**(H-I')** Electron micrographs of third instar larval wing imaginal discs from *wts-Venus* animals that also express APEX2-GBP in the posterior compartment, under the control of *en-GAL4*. A dashed orange line indicates the anterior-posterior compartment boundary in (**H' and I**). Note the elevated DAB signal at the AJs and in the basal-most region of the lateral membranes (orange arrowheads in **H-I'**). Some cells exhibited DAB signal in the sub-apical region, above the AJs (white arrows in **H''**). Elevated DAB signals were present specifically in cells of the posterior wing disc, but not in control, anterior cells. The basement membrane is marked (blue '*' in **I and I'**).

Scale bars are indicated in image panels. All confocal microscopy images are maximum intensity projections. Data is representative of at least 3 specimens per condition.

**Kroeger *et al.*, Figure S1, related to Figure 1**

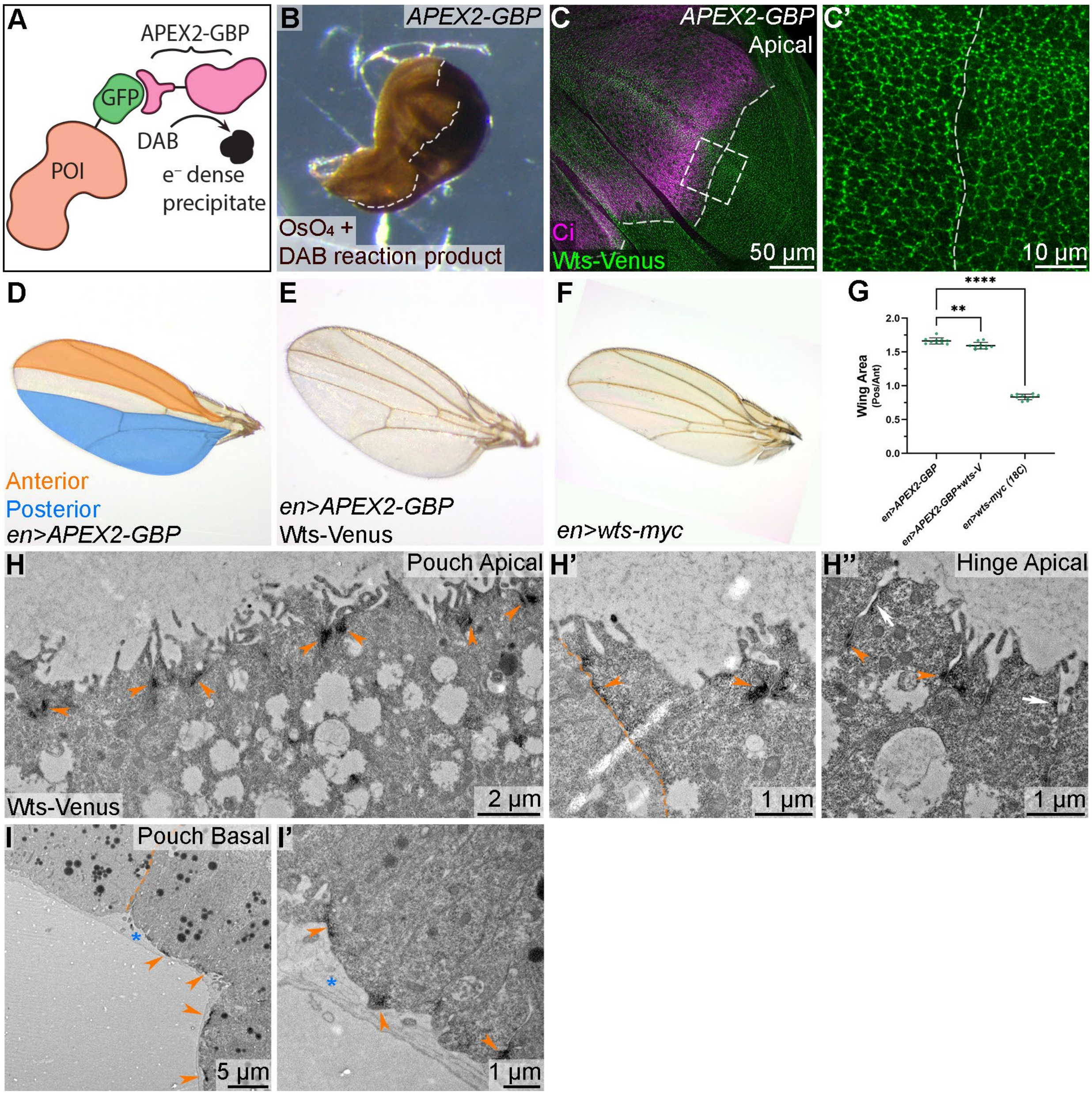

**Figure S2. Spatial analysis of basal Warts puncta in epithelial tissues.**

**(A)** Confocal microscope images of the apical region of a third instar larval wing imaginal disc. Wts-Venus is green and mRFP-NLS is magenta in the merged images, and both are greyscale in the non-merged images. Mosaic wing disc tissue is comprised of three cellular populations: *wts-Venus* homozygous clones (expressing Wts-Venus, green), *mRFP-NLS* homozygous twin-spot clones (magenta), and heterozygous cells (expressing both Wts-Venus and mRFP-NLS). Boxed region is shown at higher magnification in (**A'-A'''**). Dashed lines highlight homozygous clone boundaries.

**(B)** Confocal microscope image of the basal surface of a wing imaginal disc expressing Wts-Venus (greyscale). Boxed region (1) indicates the pouch whilst (2) indicates the ventral pouch-hinge boundary fold.

**(C)** Super-resolution Airyscan images of the apical pouch region of a third instar larval wing imaginal disc. Wts-Venus is green and E-cad-3xtRFP is magenta.

**(D and E)** Basal hinge region of wing discs expressing either GFP-Wts (green in **D-D''**) or mCitrine-Wts (greyscale in **E**). Phalloidin-TRITC (marks F-actin) is magenta in (**D and D'**). Like Wts-Venus, both of these proteins are present in basal puncta. Boxed region in (**D**) is shown at higher magnification in (**D'-D''**).

**(F)** Confocal microscope images of a third instar larval antennal disc. The mosaic tissue shown contains clonal populations of *wts-Venus* and *mRFP-NLS* cells. Boxed region is shown at higher magnification in (**F'-F'''**). Dashed lines highlight clone boundaries. At a cross section in the tissue, both AJ-associated Wts-Venus (arrows), and basal Wts-Venus puncta are indicated (arrowheads).

**(G)** Super-resolution Airyscan images of the basal region of a third instar larval eye disc expressing Wts-Venus (green) and Rhea-mCherry (blue). Boxed regions are shown at higher magnification in (**F'-F'''**). Dashed line in close up image indicates the region quantified in (**H**).

**(H)** Line profile analysis of Wts-Venus and Rhea-mCherry. The line profile (marked g-g' in **G'**), runs from left to right across the graph. Basal Wts-Venus signal peaks (marked with black arrows) do not correlate with Rhea-mCherry.

**(I)** Super-resolution Airyscan image of the basal pouch region of a third instar larval wing disc expressing Wts-Venus (green) and E-cad-3xtRFP (magenta). Dashed lines indicate the regions (1) and (2), which are quantified in (**J and J'**).

**(J)** Line profiles of Wts-Venus and E-cad-3xtRFP at basal puncta. Wts-Venus and E-cad-3xtRFP are closely spatially correlated (e.g. black arrows in **I**), but do not always show a linear relationship in fluorescence intensity (black arrowheads in **I'**).

Scale bars are indicated in image panels. All images are maximum intensity projections except for (**D-D''**, **F-F'''**, and **I**), which are single z-slices. Data is representative of at least 5 specimens per condition.

### Kroeger *et al.*, Figure S2, related to Figure 1

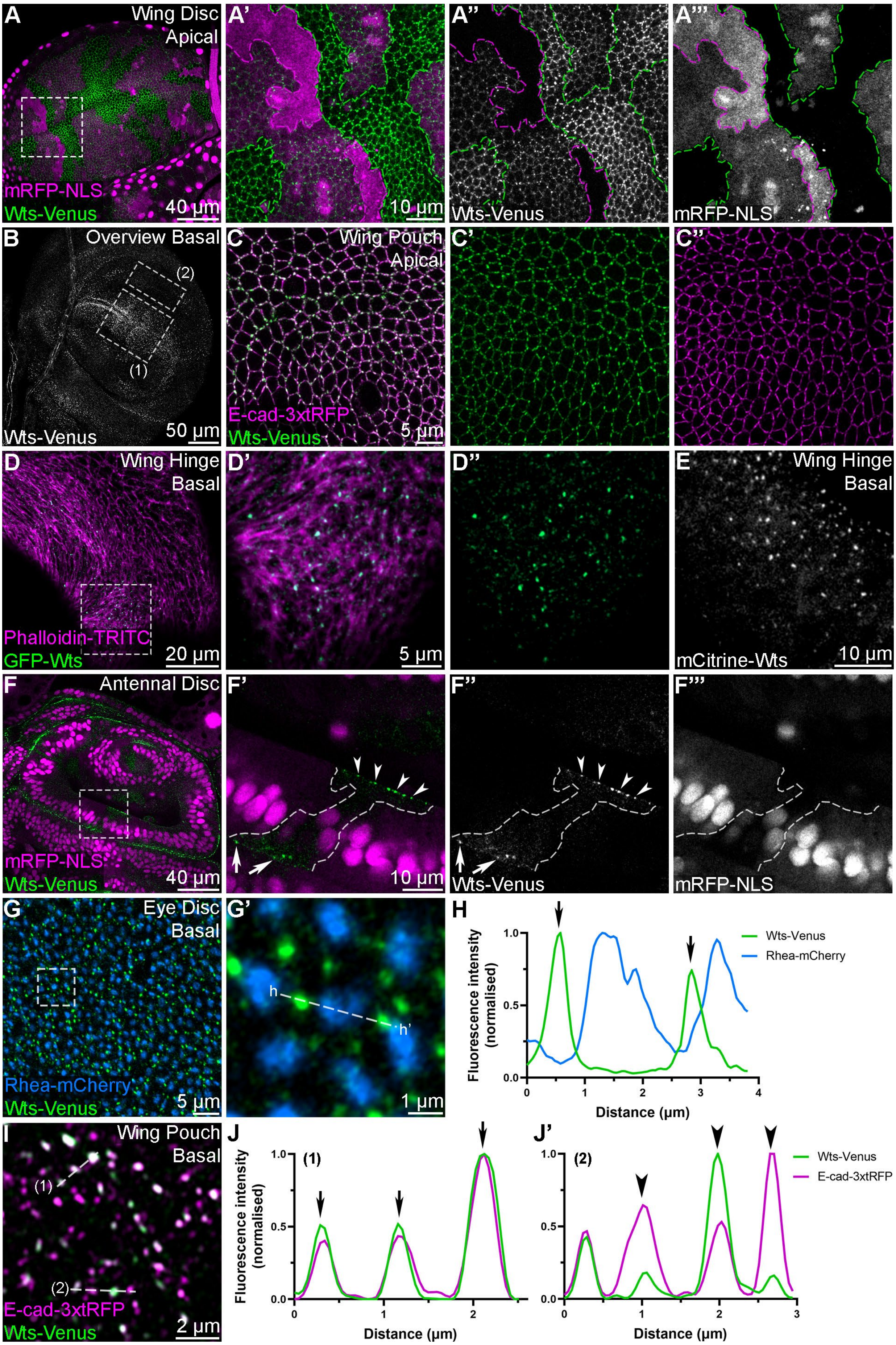

**Figure S3. Ajuba, E-cadherin and α-Catenin are required for Warts to localise at adherens junctions.**

**(A-D)** Confocal microscope images of the apical pouch region of third instar larval wing imaginal discs. Wts-Venus is green and antibody-detected Ci marks the anterior compartment (magenta) in **A-C**). Jub-GFP is in greyscale in (**D**). The indicated RNAi transgenes were expressed in the posterior compartment under the control of *en-GAL4*. Dashed lines indicate anterior-posterior compartment boundaries, posterior is to the right. Boxed regions are shown at higher magnification in **A', B'** and **C'**).

**(E)** Chart displaying the relative abundance of Jub-GFP in the posterior compared to anterior compartment of wing imaginal discs expressing the indicated RNAi lines. n = 3 for each condition. Data are represented as mean ± SD, p values were obtained using a one-way ANOVA and Dunnett's multiple comparison tests, **** = p<0.0001.

**(F-I)** Confocal microscope images of the basal hinge (**F-F'**) or apical pouch region (**G-I'**) of third instar larval wing imaginal discs. Wts-Venus is green and CD8-RFP (magenta) marks clones expressing the indicated RNAi transgenes for 48 h. (**G-I**) are higher magnification images than (**F**). Scale bars are indicated in image panels. All images are maximum intensity projections. Data is representative of between 3-10 specimens per condition.

### Kroeger *et al*., Figure S3, related to Figure 3

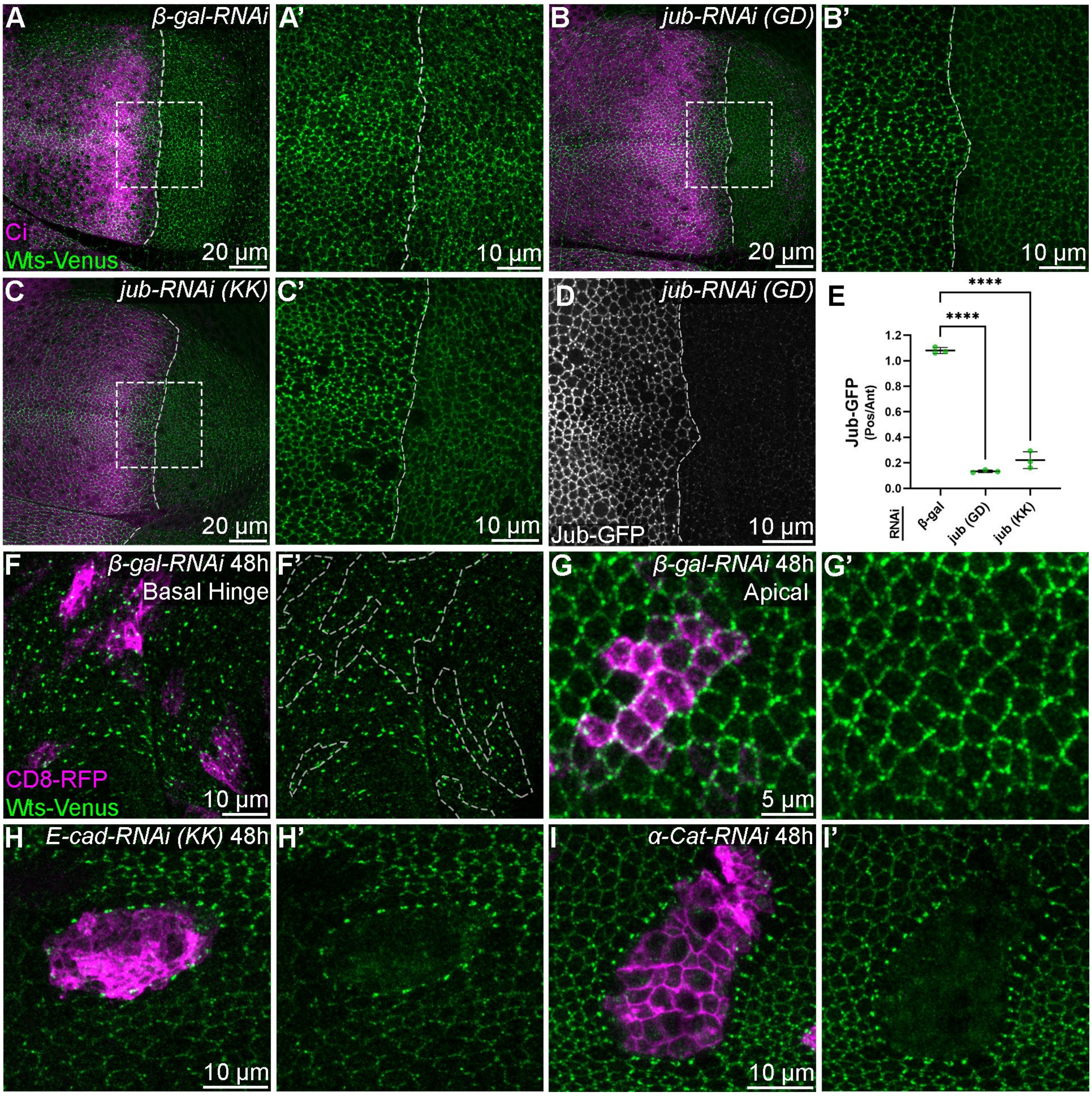

**Figure S4. Most Hippo pathway proteins are enriched at apical, but not basal, membranes of epithelial cells.**

**(A-G)** Super-resolution Airyscan images (**A, A'**) and confocal microscope images (**B-G'**) of third instar larval wing imaginal discs expressing different tagged Hippo pathway proteins. (**A-E', G)** and (**G'**) display planar images of the apical domain of wing imaginal discs, and (**F**) and (**F')** are cross-section images. In (**F)** and (**F')**, dashed grey lines indicate basal surfaces (also marked with arrowheads), whilst apical membrane regions are indicated by arrows. In (**A, A'**) Hpo-Venus is orange and mRFP-NLS is magenta. Mosaic wing disc tissue in (**A**) is comprised of *hpo-Venus* homozygous clones, *mRFP-NLS* homozygous twin-spot clones, and heterozygous cells (which express both). GFP-Mats is blue (**B, B'**), Sav-Venus is cyan (**C, C'**) and Mer-Venus is yellow (**D, D'**). Kibra-Venus is in greyscale in (**E-F'**) and mRFP-NLS is magenta in (**E and F**). Mosaic wing disc tissues in (**E-F'**) are comprised of three cellular populations: *kibra-Venus* homozygous clones, *mRFP-NLS* homozygous twin-spot clones (marked with magenta dashed lines in **E'** and **F'**), and heterozygous cells (which express both). Ex-eYFP is red in (**G** and **G'**). Boxed regions are shown at higher magnification. Wing discs are live in (**C-D'**) and fixed in all other images. Scale bars are indicated. All images are maximum intensity projections except for (**A-A'** and **F-F'**), which are single z-slices. Data is representative of at least 5 specimens per condition.

### Kroeger *et al.*, Figure S4, related to Figure 4

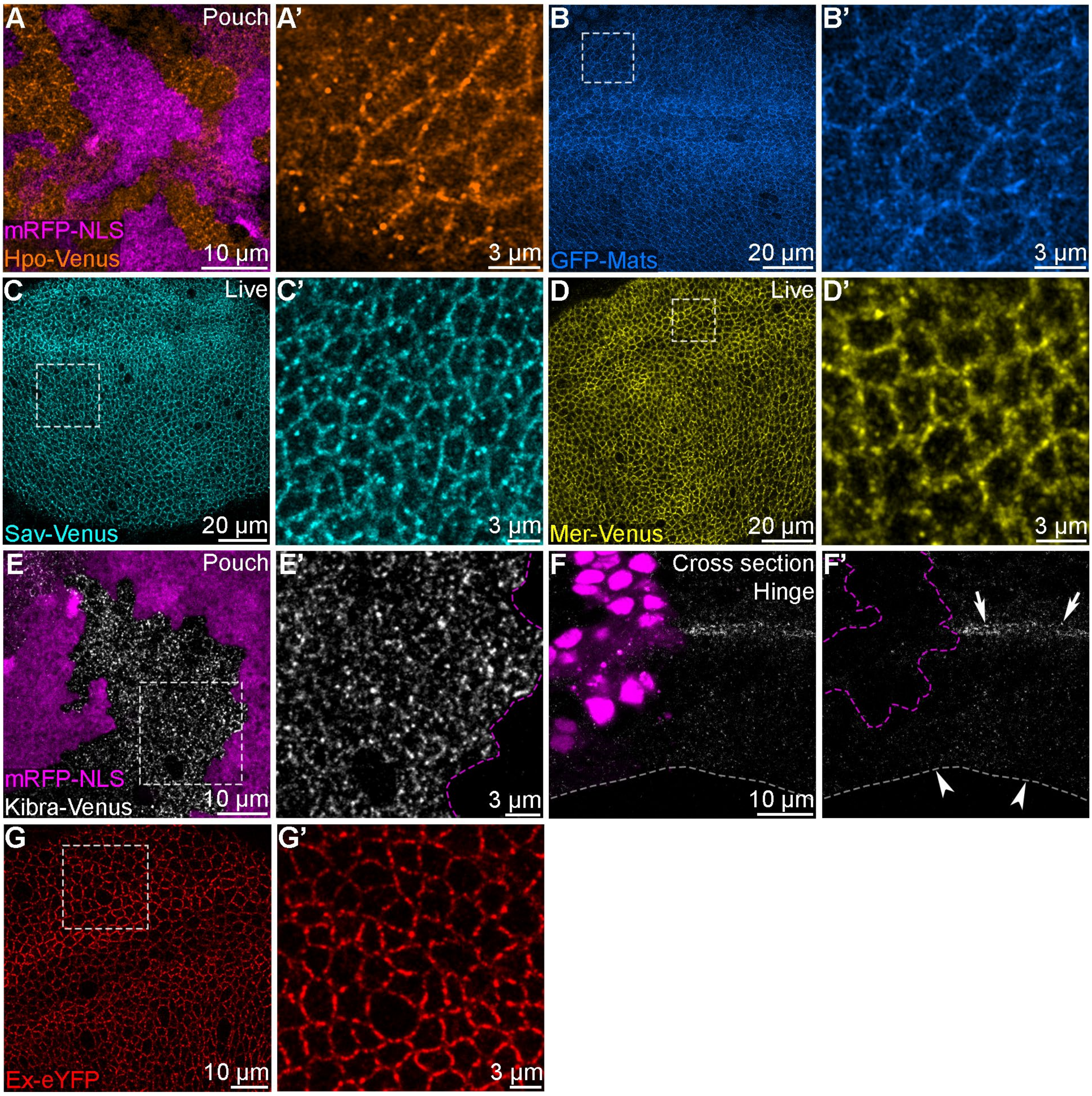

**Figure S5. Warts localises to basal spot junctions in the pupal notum.**

**(A and B)** Confocal microscope images of basal or apical regions of the pupal notum, captured at the indicated times after puparium formation (hAPF). The tissue is comprised of three cellular populations: *wts-Venus* homozygous clones (expressing Wts-Venus, green or in grayscale), *mRFP-NLS* homozygous twin-spot clones (magenta or in grayscale), and heterozygous cells (expressing both Wts-Venus and mRFP-NLS). Dashed lines highlight clone boundaries. Boxed regions are shown at higher magnification in (**A'-A'''** and **B'-B'''**).

**(C-F)** Confocal microscope images of basal regions of the pupal notum at the indicated times after puparium formation. Wts-Venus is green in (**C and E**), E-cad-3xtRFP is magenta in (**C and D**), Jub-GFP is green in (**D**), Rhea-mCherry is blue in (**E**), MyoII-3xmKate2 is magenta in (**F**), and Rhea-EGFP is cyan in (**F**). Punctate Wts-Venus and Jub-GFP fluorescent signals are closely correlated spatially with E-cad-3xtRFP (arrowheads in **C-D''**). Boxed region in (**E**) is shown at higher magnification in (**E'-E'''**). Dashed line in (**F**) indicates the region quantified in (**G**).

**(G)** Line profile analysis of MyoII-3xmKate2 and Rhea-EGFP. The line profile (g-g' in **F**), runs from left to right across the graph.

Scale bars are indicated in image panels. All images are maximum intensity projections except for (**C-D''**), which are single z-slices. Data is representative of 5-10 specimens per condition.

### Kroeger *et al*., Figure S5, related to Figure 5

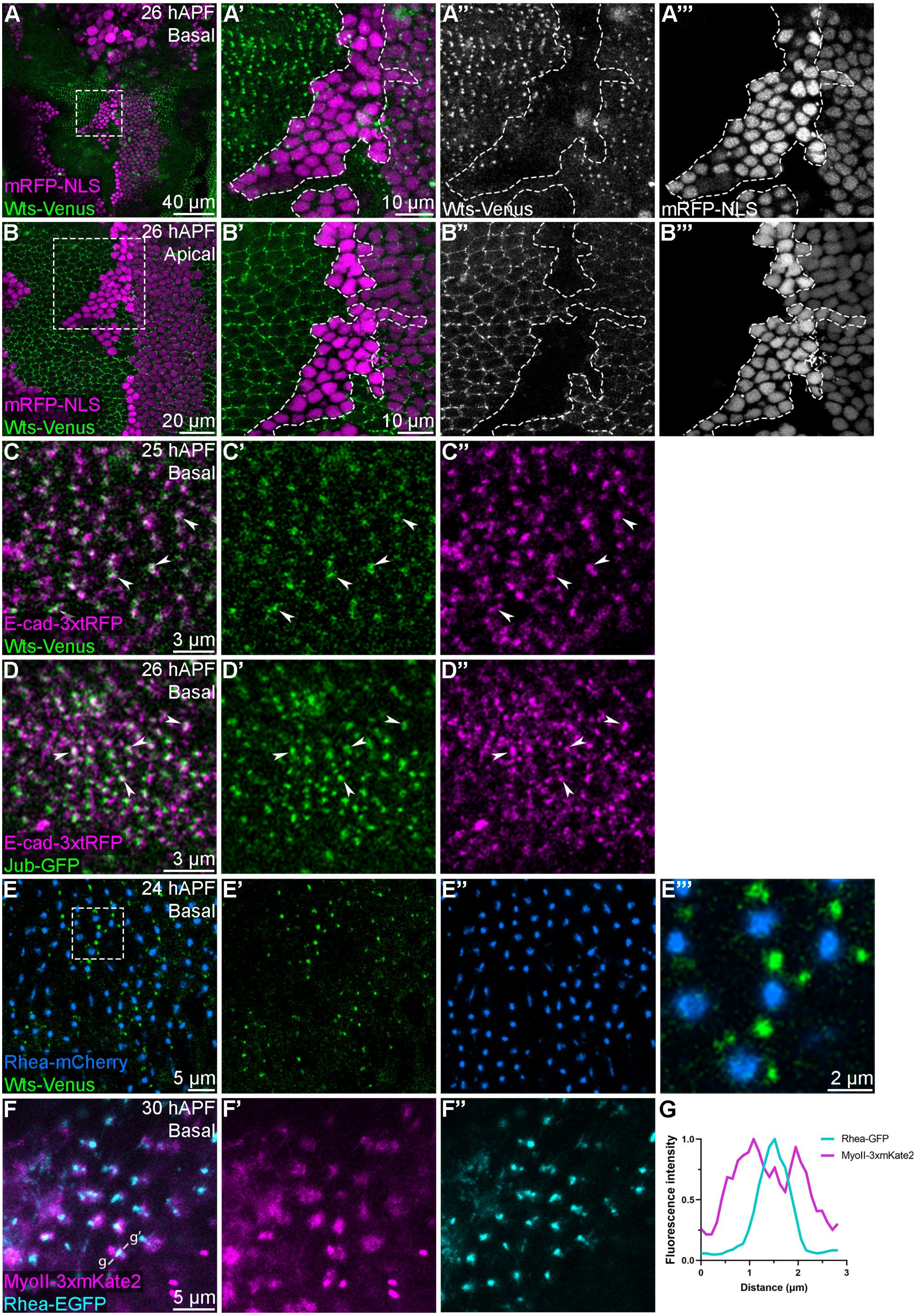

**Figure S6. Warts localises to basal spot junctions in wing imaginal discs and the pupal notum in a spatially restricted manner.**

**(A-D)** Confocal microscope images of the basal region of third instar larval wing imaginal discs at 0 h (**A, A', E**), 12 h (**B, B', F**), 24 h (**C, C', G**) and 48 h (**D, D', H**) after third instar ecdysis. Wts-Venus is green, MyoII-3xmKate2 and E-cad-3xtRFP are magenta. Boxed regions in (**A-D**) are shown at higher magnification in (**A-D')** and these are super-resolution Airyscan images.

**(I-L)** Confocal microscope images of the basal region of the central-posterior region of the pupal notum epithelium at 17 h (**I-I''**), 18 h (**J-J''**), 24.5 h (**K-K''**) and 29.5 h (**L-L''**) after puparium formation. Wts-Venus is green and Rhea-mCherry is blue. Images **(I-L**'') are projections with z-planes manually cropped to remove autofluorescent regions and to optimally display the basal surface of the epithelium. Boxed regions in (**I'-L'**) are shown at higher magnification in (**I''-L''**). Scale bars are indicated in image panels. All wing disc images are single z-slices except for (**C** and **F**), which are maximum intensity projections. Data is representative of 3-10 specimens per condition.

### Kroeger *et al.*, Figure S6, related to Figure 5

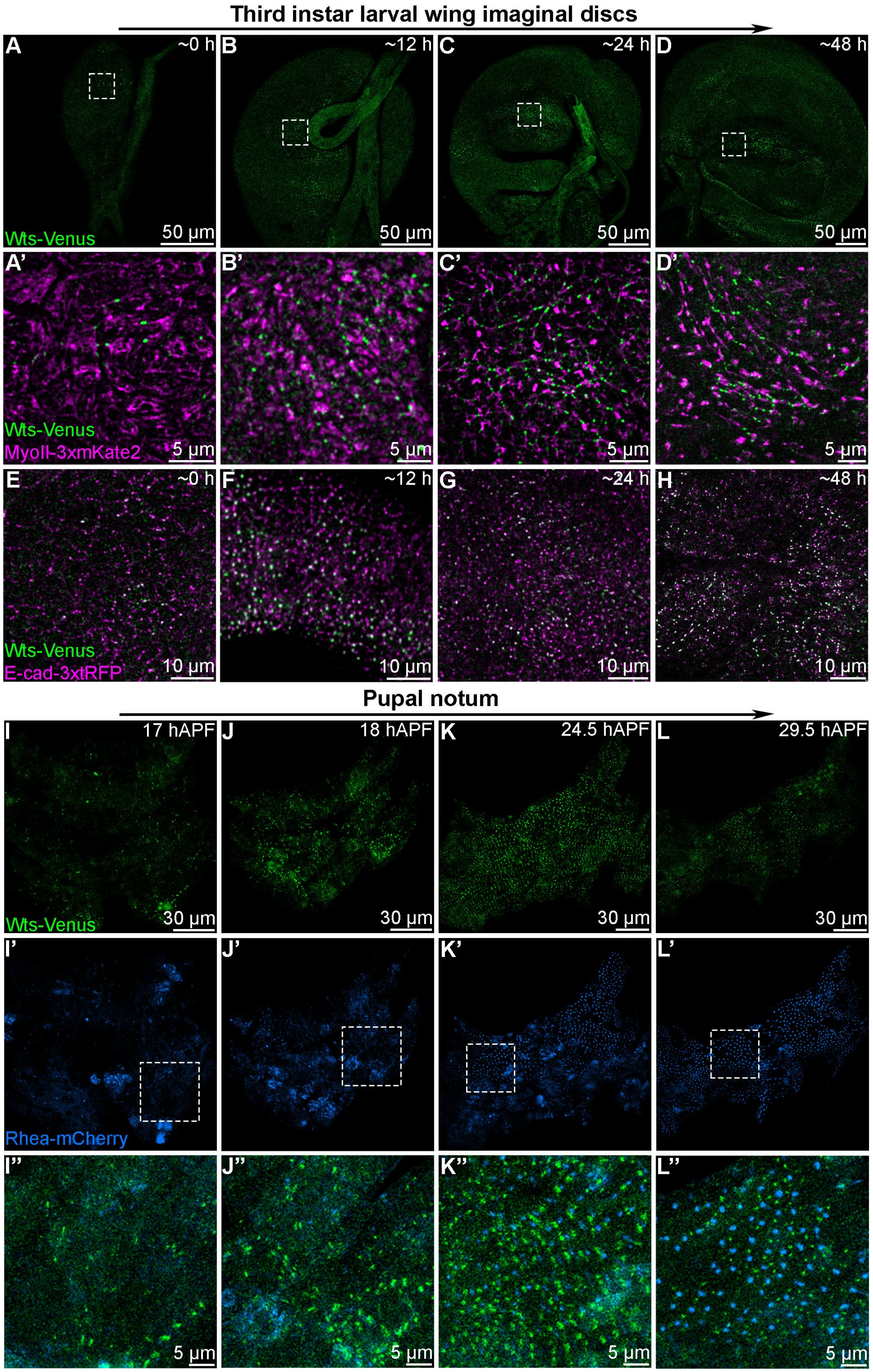

**Figure S7. Focal adhesions influence Warts recruitment to basal spot junctions, and Yorkie subcellular localization and activity.**

**(A-D)** Confocal microscope images of the basal pouch region of third instar larval wing imaginal discs. Rhea-mCherry is in greyscale in (**A-B**), antibody-detected Mys is greyscale in (**C-D**). The indicated RNAi transgenes were expressed for 48 h in the posterior compartment under the control of *en-GAL4*. Dashed lines indicate the anterior-posterior boundary.

**(E-F)** Charts displaying the relative abundance of Rhea-mCherry (**E**) and Mys (**F**) in the posterior compared to anterior compartment of wing imaginal discs expressing the indicated RNAi lines. n = 3, 3, 4, 2 in (**E**) and 2 each in (**F**). Data are represented as mean ± SD, p values in (**E**) were obtained using a one-way ANOVA and Dunnett's multiple comparison tests, **** = p<0.0001.

(**H-N'**) Confocal microscope images of third instar larval wing imaginal discs. Images of the basal plane of the ventral pouch-hinge boundary fold in are shown in (**G-J'**), and the central plane of the same region in (**K-L**), whilst the apical plane of the pouch is shown in (**M-N'**). The indicated rhea-RNAi transgenes were expressed for 48 h in the posterior compartment under the control of *en-GAL4*. Dashed lines indicate the anterior-posterior boundary in (**G-L**). E-cad-3xtRFP is magenta and Wts-Venus is green in (**G-J'**), Yki is in greyscale in (**K**), β-gal driven by *ex-lacZ* is in greyscale in (**L**), Hpo-Venus is orange and Arm is blue in (**M-N'**).

Scale bars are indicated in image panels. All images are maximum intensity projections except for (**C-D and M-N**), which are single z-slices. Data is representative of 2-10 specimens per condition.

### Kroeger *et al.*, Figure S7, related to Figure 7

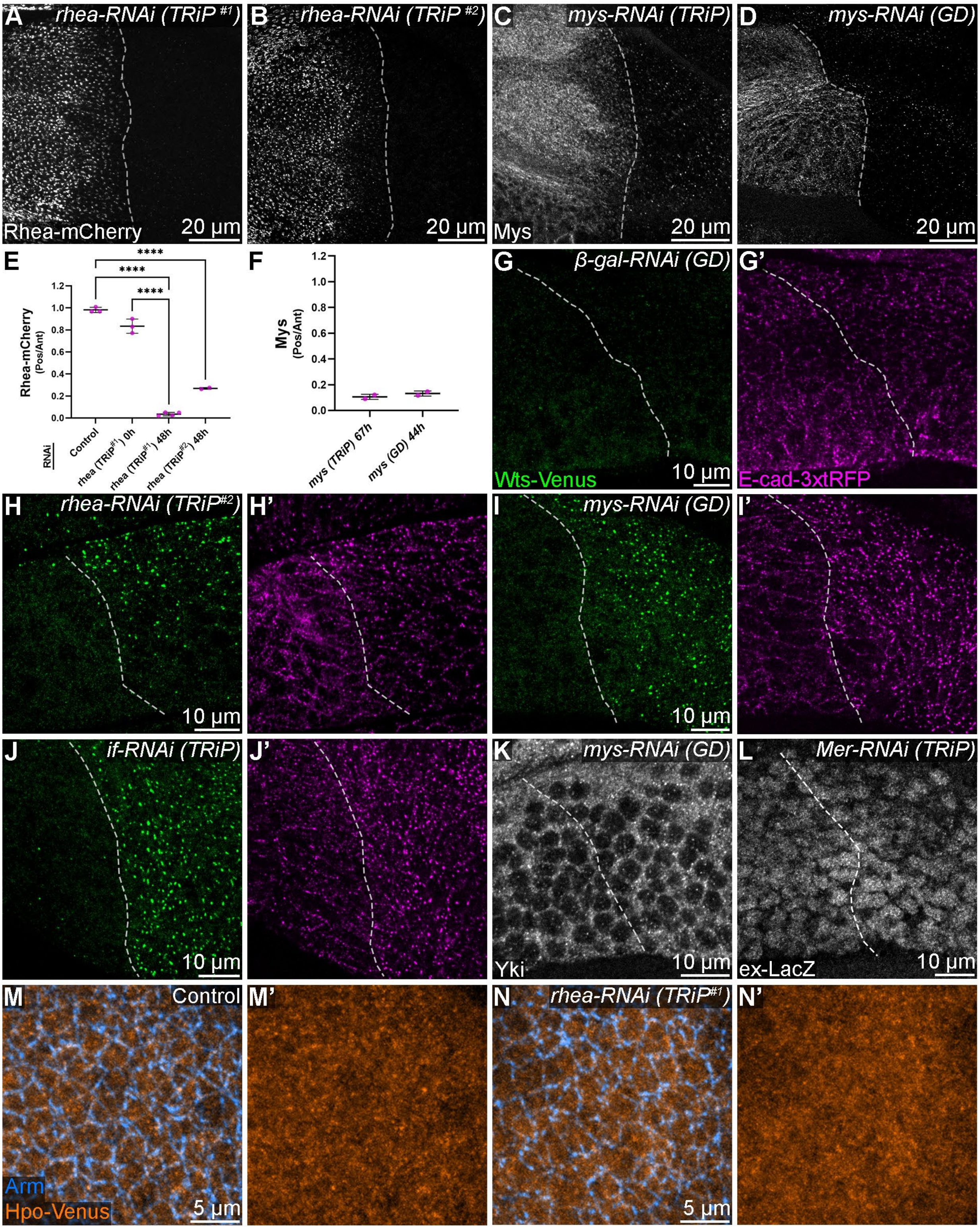

**Figure S8. Genetic depletion of focal adhesion proteins moderately influences apical abundance of Warts protein.**

**(A-A''')** Confocal microscope images of the basal regions of third instar larval wing imaginal discs. Rhea-EGFP is in grayscale, the boxed regions in (**A**) are labelled (1-3) and are magnified in (**A'-A'''**). Images (**A'-A''**) are super-resolution Airyscan acquisitions.

**(B-E''')** Confocal microscope images of the apical regions of third instar larval wing imaginal discs. E-cad-3xtRFP is magenta, Wts-Venus is green and antibody-detected Ci marks the anterior compartment (yellow). The indicated RNAi transgenes were expressed for 48 h or 72 h in the posterior compartment under the control of *en-GAL4*. Dashed lines indicate the anterior-posterior boundary, posterior is to the right. White arrowheads indicate morphological disruptions in small regions of the apical domain. Boxed regions are shown at higher magnification in (**B'''**, **C'''**, **D'''** and **E'''**).

**(F-H)** Charts displaying the relative amount of apical junctional Wts (**F**) or total Wts (**G** and **H**) present in the posterior compared to anterior compartment of wing imaginal discs expressing the indicated RNAi lines. In (**F**) n = 4, 4, 4, 3; (**G**) n = 2, 2, 3; (**H**) n = 4, 4, 2. Data are represented as mean ± SD, p values were obtained using a one-way ANOVA and Dunnett's multiple comparison tests, ns = not significant, ** = p<0.01, *** = p<0.001.

Scale bars are indicated in image panels. All images are maximum intensity projections. Data is representative of 3-5 specimens per condition.

**Kroeger *et al.*, Figure S8, related to Figure 7**

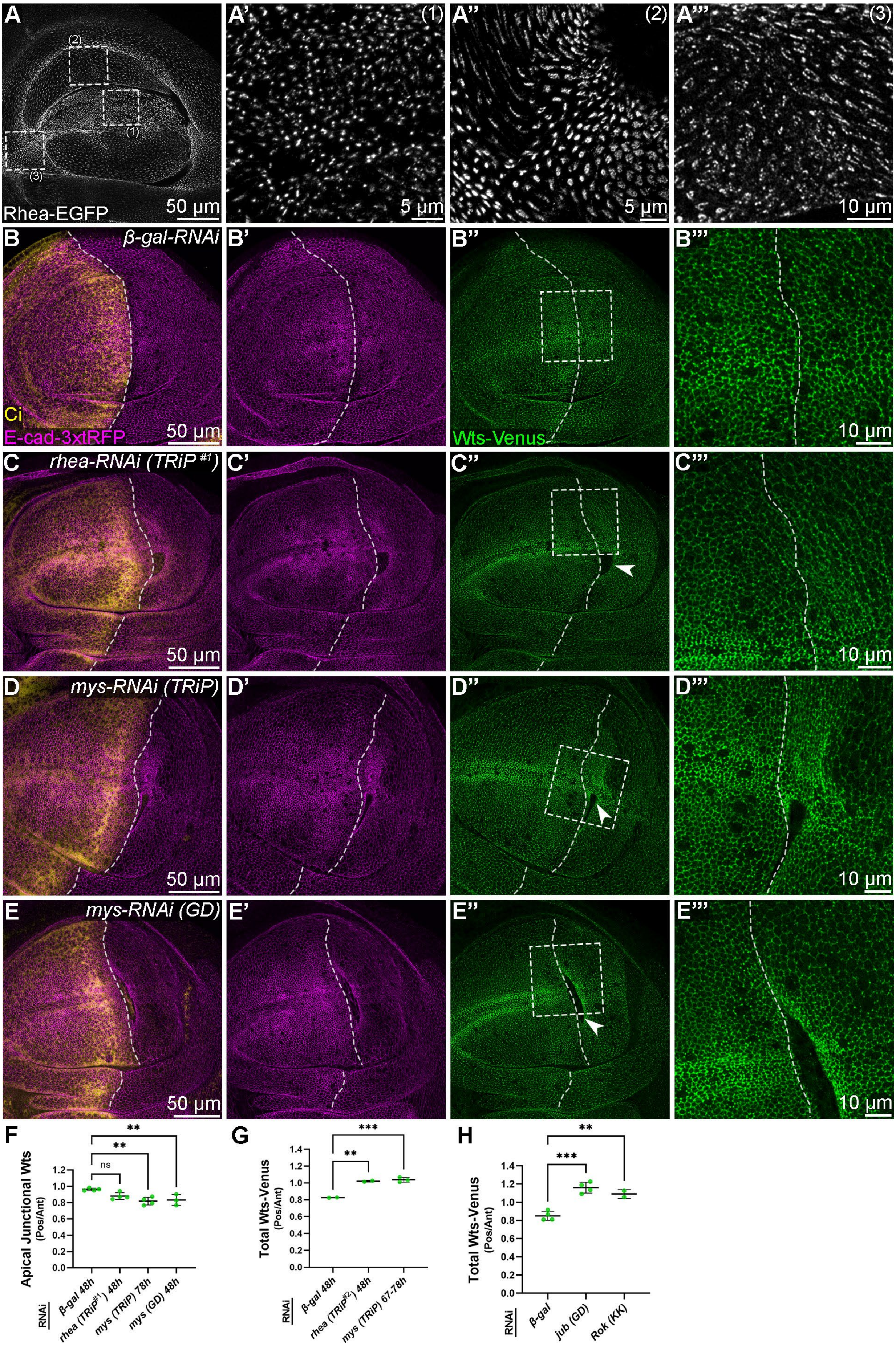

**Figure S9. Forced re-localization of Warts to basolateral or apical membranes modulates its ability to control Yorkie activity and wing size.**

**(A-E')** Confocal microscope images of third instar larval wing imaginal discs. Wts-Venus is green, mCherry is magenta, E-cad-3xtRFP is blue. Dashed lines indicate the anterior-posterior boundary, posterior is to the right. Planar images of the wing pouch are shown in (**A-B'**) at the apical (**A, A', D, D'**) and basal (**B, B', E, E'**) surfaces. The boxed regions in (**B** and **E**) are magnified in (**B'** and **E'**). Images of the apical cell junctions in a fold between the hinge and pouch regions are shown in (**C-C'''**). Arrows indicate Wts-Venus at AJs, arrowheads indicate Wts-Venus that is apical of the AJs.

**(F-F''')** Adult female *D. melanogaster* wings expressing the indicated transgenes under the control of *en-GAL4*.

**(G)** Chart showing the ratio of posterior wing area compared to anterior wing area for the indicated genotypes. n=9, 8, 10, 7, 6, 10, 9, 10. Data are represented as mean ± SD, p values were obtained using a one-way ANOVA and Šídák's multiple comparisons tests, ns = not significant, **** = $p<0.0001$.

(**H-J**) Confocal microscope images of the ventral pouch-hinge boundary fold of third instar larval wing imaginal discs, transcriptional activity of the *ex* gene was detected by β-gal antibodies (greyscale). The indicated transgenes were expressed in the posterior compartment under the control of *en-GAL4*. Dashed lines indicate the anterior-posterior boundary, posterior is to the right.

**(K)** Chart comparing the ratio of β-gal in posterior compared to anterior wing discs expressing the indicated transgenes. n=7, 4, 5, 5. The dotted line indicates the mean ex-LacZ measurement from *mCherry-RNAi* samples. Data are represented as mean ± SD, p values were obtained using a one-way ANOVA and Dunnett's multiple comparison tests, *** = $p<0.001$, **** = $p<0.0001$.

Scale bars are indicated in image panels. All confocal images are maximum intensity projections except (**C-C'''** and **H-J**), which are single z-slices. Data is representative of 5-10 specimens per condition.

### Kroeger *et al.*, Figure S9, related to Figure 7

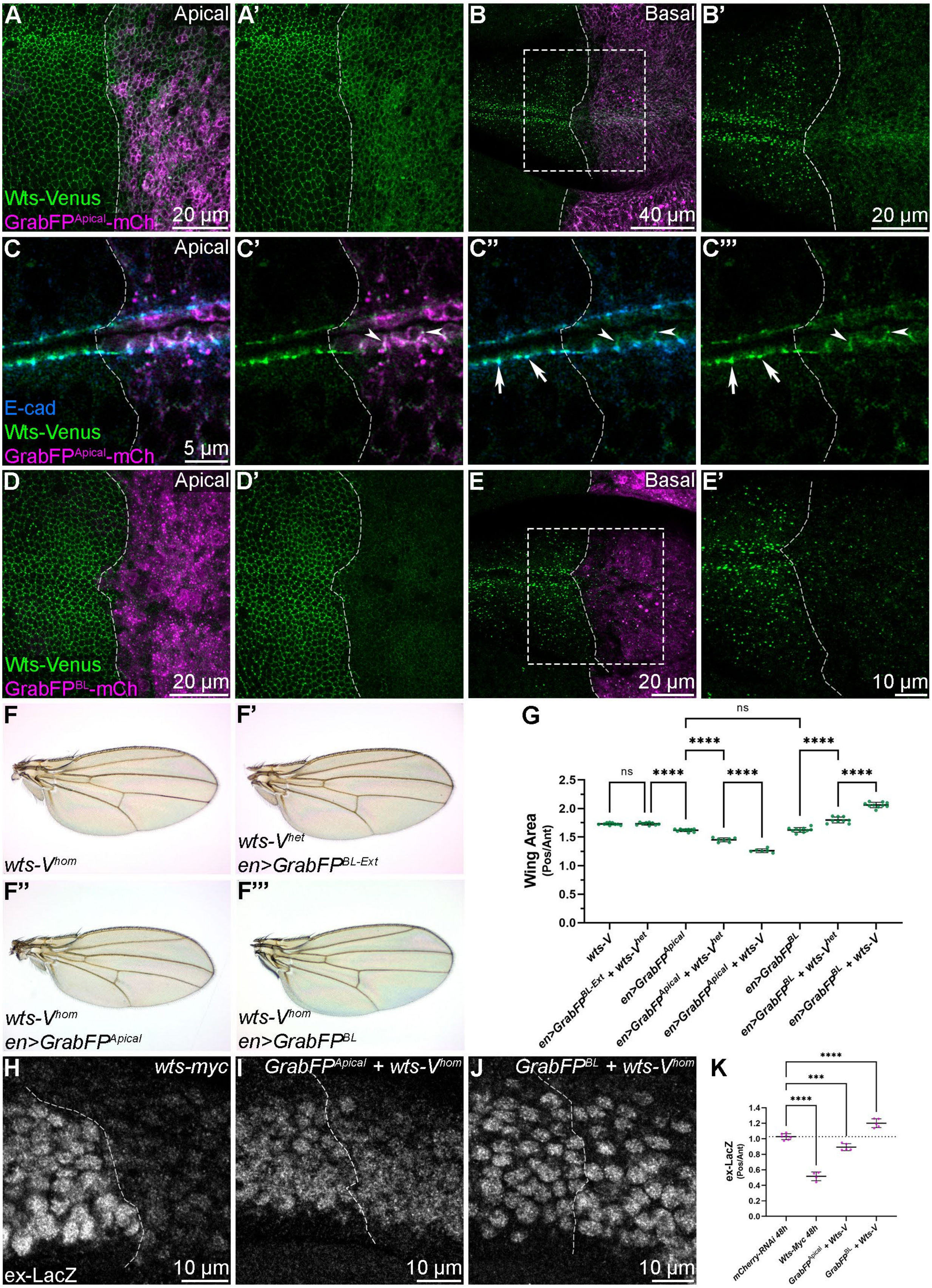

**Video S1. Three-dimensional reconstruction of Warts-Venus localization in a larval wing imaginal disc.**

Video displaying a confocal microscope 3D-reconstruction of the wing pouch region of a third instar larval wing imaginal disc. Wts-Venus is green and CD8-mRFP is magenta. The epithelium is oriented at the start of the video with the apical side at the top and basal side of the bottom.

**Video S2. Fluorescence recovery after photobleaching experiments on Warts-Venus at AJs and basal spot junctions.**

Example FRAP videos acquired at the apical (left) or basal (right) surface of the pupal thorax epithelium between 21-24 h APF. Wts-Venus signal within the yellow ROI was photobleached using a high intensity 514 nm laser pulse after 5 frames (10 sec) at the level of the adherens junctions (left) or basal spot junctions (right), and fluorescence recovery was measured over the subsequent 2 min 50 sec. ROIs used for background subtraction (blue) and bleach correction (grey) are shown.

**Video S3. Live-cell dynamics of basal Warts and Myosin in the pupal notum.**

Live time-lapse confocal microscopy of the posterior midline region of the pupal notum epithelia at 21 hAPF captured at 1 min intervals for 1 h. Wts-Venus is green and MyoII-3xmKate2 is magenta. Topmost portion of the video includes autofluorescent cells and structures in the body cavity, underlying the epithelium. Scale bar is indicated. Video frame rate is 2fps.

**Video S4. Recruitment of Warts to basal spot junctions is coincident with enrichment of basal-medial actomyosin.**

Live time-lapse confocal microscopy of a basal region of the pupal notum epithelia at 25 hAPF captured at 1 min intervals for 8 mins. Wts-Venus is green and MyoII-3xmKate2 is magenta. Note the dynamic changes in Wts-Venus and MyoII-3xmKate2 at the basal region of the indicated cell (arrows). Scale bar is indicated. Video frame rate is 2fps.
